# Cell type-specific enhancer-promoter connectivity maps in the human brain and disease risk association

**DOI:** 10.1101/778183

**Authors:** Alexi Nott, Inge R. Holtman, Nicole G. Coufal, Johannes C.M. Schlachetzki, Miao Yu, Rong Hu, Claudia Z. Han, Monique Pena, Jiayang Xiao, Yin Wu, Zahara Keuelen, Martina P. Pasillas, Carolyn O’Connor, Simon T. Schafer, Zeyang Shen, Robert A. Rissman, James B. Brewer, David Gosselin, David D. Gonda, Michael L. Levy, Michael G. Rosenfeld, Graham McVicker, Fred H. Gage, Bing Ren, Christopher K. Glass

## Abstract

Unique cell type-specific patterns of activated enhancers can be leveraged to interpret non-coding genetic variation associated with complex traits and diseases such as neurological and psychiatric disorders. Here, we have defined active promoters and enhancers for major cell types of the human brain. Whereas psychiatric disorders were primarily associated with regulatory regions in neurons, idiopathic Alzheimer’s disease (AD) variants were largely confined to microglia enhancers. Interactome maps connecting GWAS variants in cell type-specific enhancers to gene promoters revealed an extended microglia gene network in AD. Deletion of a microglia-specific enhancer harboring AD-risk variants ablated *BIN1* expression in microglia but not in neurons or astrocytes. These findings revise and expand the genes likely to be influenced by non-coding variants in AD and suggest the probable brain cell types in which they function.

**One Sentence Summary:** Identification of cell type-specific regulatory elements in the human brain enables interpretation of non-coding GWAS risk variants.

## Introduction

The central nervous system is an extraordinarily complex organ consisting of a multitude of diverse and highly interconnected cells. Recent advances in single cell sequencing technologies have dramatically advanced our appreciation of the molecular phenotypes of neurons, microglia, astrocytes, oligodendrocytes and other cell types that reside within the brain (*1–4*). In contrast, transcriptional mechanisms that control their developmental and functional properties in health and disease remain less well understood. Studies in animal models have provided deep insights into conserved processes, but significant differences between species often limit the translatability of findings in model systems to humans (*5*).

Genome-wide association studies (GWASs) provide a genetic approach to identifying molecular pathways involved in complex traits and diseases by defining associations between genetic variants and phenotypes of interest (*6, 7*). Large-scale GWASs have discovered hundreds of single nucleotide polymorphisms (SNPs) that are associated with the risk of neurological and psychiatric disorders. The vast majority of disease-risk genetic variants are located in non-coding regions of the genome (*7*) and the specific cell type(s) in which the disease-risk variants are active is often unclear. Furthermore, linkage-disequilibrium between disease-risk variants at GWAS significant loci complicates the identification of causal variants. Collectively, these limitations have hindered the assignment and interpretation of disease-risk genes.

GWAS-identified risk variants that are located in non-coding regions of the genome can exert phenotypic effects through several mechanisms, including perturbation of transcriptional enhancers and promoters (*6*). While promoters provide the essential sites of transcriptional initiation of mRNAs, they are frequently not sufficient to direct appropriate developmental and signal-dependent levels of gene expression (*8, 9*). This additional information is provided by enhancers, short regions of DNA that, when bound by transcription factors, enhance mRNA expression from target promoters. Enhancers can reside hundreds of thousands of base pairs away from their target gene and their function is generally considered to depend on three-dimensional enhancer-promoter interactions (*10*). Enhancer selection is driven by cell type-specific combinations of lineage-determining transcription factors that, in turn, specify the binding of signal-dependent transcription factors. As a consequence, each cell has a unique enhancer repertoire that underlies its particular pattern of gene expression and enables cell type-specific responses to intra- and extra-cellular signals (*11–13*). By defining the enhancer landscape of a cell, it is possible to infer key transcription factors and environmental signals guiding that cell’s identity and functional potential. Accordingly, genetic variation affecting enhancer selection and function is considered to be a major determinant of differences in cell type-specific gene expression between individuals (*14*).

Despite their importance, the enhancer landscapes of different brain cell types remain largely unknown. We recently reported transcriptomes and enhancer maps of human microglia isolated from surgically resected tissue (*15*). We found that AD-risk genes are highly expressed in microglia compared to whole cortex. However, interpretation has been limited by a lack of knowledge of enhancer landscapes in other relevant cell types. Furthermore, recent studies in AD have relied on bulk tissue (*16–18*), potentially masking epigenomic changes that occur in rare cell types. For example, microglia constitute around 5% of brain cells, therefore analysis of bulk tissue renders microglia-specific enhancers difficult to detect. Emerging single nuclei ATAC-seq methods provide a powerful method to define cell type-specific patterns of open chromatin corresponding to enhancers and promoters (*2, 19*). However, these methods cannot distinguish between active and poised regulatory elements. Importantly, previous studies of gene regulatory regions in the brain do not provide information on the connectivity of putative enhancers with their target promoters.

To advance our knowledge of enhancer connectivity in the brain, we established protocols for cell type-specific nuclei sorting using frozen human brain tissue. Using these methods, we generated high-quality atlases of putative active enhancers and promoters in microglia, neurons, oligodendrocytes and astrocytes and three-dimensional connectivity maps for microglia, neurons, and oligodendrocytes. Genetic variants associated with complex brain traits and diseases exhibit strikingly different patterns of enrichment in cell type-specific enhancers. In combination with connectivity maps, we provide evidence for microglia-specific targets for a subset of AD GWAS variants, reassign a subset of variants to different targets than the closest gene, and expand assignments of target genes for others. Lastly, using in vitro differentiation of human pluripotent stems cells, we validate the microglia-specific function of a putative microglia enhancer harboring an AD variant that is connected to the BIN1 promoter.

## Results

### Promoter and enhancer atlases of microglia, neurons, astrocytes and oligodendrocytes

To characterize transcriptional regulatory elements within different cell types of the human brain, we established a cell type-specific nuclei isolation protocol for frozen tissue (Fig. 1A). Nuclei were isolated from cortical brain tissue resections taken from epilepsy patients (Table S1). Consistent with our previous methods, resected tissue excluded the brain area displaying the strongest epileptogenic activity and was determined to be histologically normal (*15*). Nuclei were immuno-labeled with the following cell type-specific markers: PU.1 for microglia, NEUN for neurons and OLIG2 for oligodendrocytes (Fig. 1B) and then isolated by florescence activated nuclear sorting (FANS). Currently, there is a lack of antibodies for nuclear astrocytic markers. However, LIM/Homeobox 2 (LHX2) is highly expressed in astrocytes and moderately expressed in neurons (*20, 21*). Using a combinatorial flow cytometry strategy (NEUN-negative, LHX2-positive), we were able to isolate astrocyte nuclei (Fig. 1B).

**Fig. 1.**
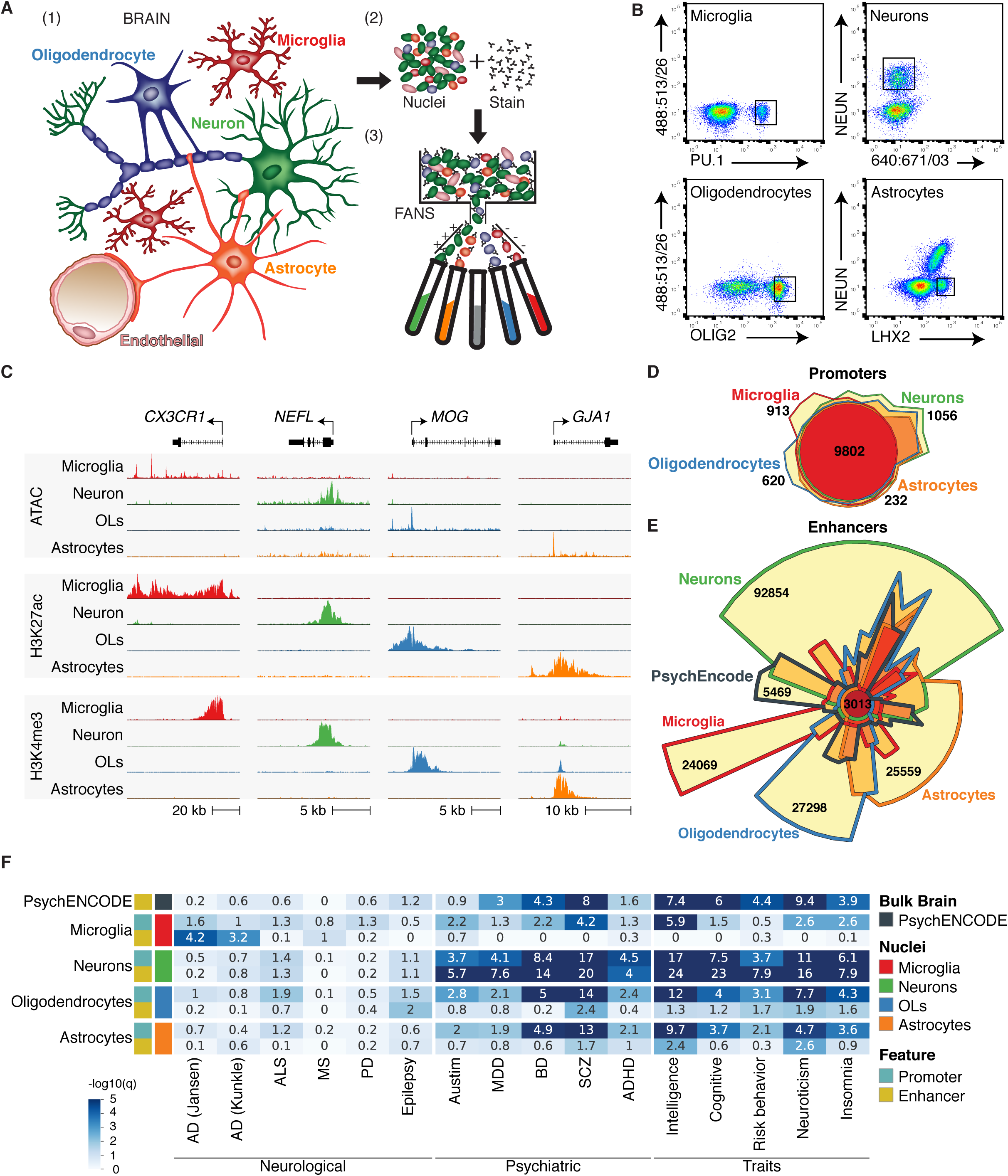
Cell type-specific genomic regulatory regions of microglia, neurons, astrocytes and oligodendrocytes are differentially enriched for GWAS risk variants for neurological and psychiatric disorders and behavioral traits. (A) Nuclei isolation strategy for generating cell type-specific datasets of the human brain. (B) Microglia, neuronal and oligodendrocyte populations were defined as PU.1^+^, NEUN^+^ and OLIG2^+^ nuclei, respectively. The astrocyte population was defined as NEUN^neg^ LHX2^+^ nuclei. (C) UCSC browser visualization of open chromatin regions defined by ATAC-seq signal (top panel), H3K4me3 abundance at promoters (middle panel) and H3K27ac at gene regulatory regions (bottom panel) for brain nuclei populations. Displayed is a representative gene for microglia (*CX3CR1*), neurons (*NEFL*), oligodendrocytes (*MOG*) and astrocytes (*GJA1*). (D) Chow-Ruskey plot of promoter regions defined for brain cell populations. (E) Chow-Ruskey plot of enhancer regions defined for cell types of the brain and enhancers defined by PsychENCODE using bulk brain. (F) Heatmap of LDSC analysis for genetic variants associated with neurological and psychiatric disorders and behavior traits displayed as -log10(q) value for significance of enrichment for promoter and enhancer regions of brain nuclei populations. Top row shows LDSC analysis for enhancer regions defined by PsychENCODE using bulk brain.

Isolated nuclei were subjected to ATAC-seq, which identifies open regions of chromatin (*22*), and H3K27ac and H3K4me3 ChIP-seq, which predominantly identify active chromatin regions and promoters, respectively (*23, 24*). ATAC-seq, and H3K27ac and H3K4me3 ChIP-seq datasets were processed according to ENCODE guidelines (see methods and Table S2) and clustered according to cell type of origin (fig. S1A-C). ATAC, H3K27ac and H3K4me3 profiles exhibited highly cell type-specific patterns (fig. S1A-C), exemplified by the cell type-specific genes *CX3CR1*, *NEFL*, *MOG* and *GJA1* (Fig. 1C). Importantly, ATAC-seq and H3K27ac datasets generated from PU.1 nuclei closely resembled those of ex vivo live microglia isolated at 98% purity using comparable resected tissue, indicating that PU.1 nuclei from frozen tissue closely recapitulated microglia in the brain (*15*) (fig. S1A, B).

To test whether epigenomic data could be used as a proxy for gene expression, we correlated ATAC-seq, H3K27ac and H3K4me3 ChIP-seq signals at promoters to published gene expression data that exist for cell types of the brain (*21*). While each of the nuclei epigenomic datasets correlated with gene expression, promoter H3K27ac most closely mirrored gene expression of brain cell types (fig. S1D). Similarly, promoters associated with signature gene sets defined for microglia, neurons, oligodendrocytes and astrocytes by PsychENCODE (*25, 26*) exhibited corresponding preferential H3K27ac (fig. S1E). OLIG2 nuclei had increased H3K27ac at the promoters of oligodendrocyte signature genes compared to oligodendrocyte precursor cell (OPC) signature genes, indicating a low number of OPCs. Comparison of NEUN promoter H3K27ac across neuronal subtype gene signatures shows an absence of differences between neuronal subtype markers, suggesting that NEUN nuclei represent a mix of excitatory and inhibitory neurons (fig. S1E).

Cell type-specific promoter activity states were then defined using differential analysis of promoter H3K27ac for each cell type compared to the average of the other cell types (fig. S2A, Table S3). H3K27ac-defined promoter activity exhibited a similar cell type-specific pattern as seen for gene expression (*21*), as well as ATAC-seq and H3K4me3 enrichment (fig. S2B-C). These H3K27ac-defined brain cell type signature genes were associated with gene ontologies representative of the cell type of origin (fig. S2D). Taken together, these analyses indicate that we were able to collect high quality epigenomic data of neurons, astrocytes, oligodendrocytes and microglia using frozen human resected tissue.

Gene regulatory regions vary according to cell of origin, yet studies identifying active promoters and enhancers of most human brain cell types in vivo are lacking. We defined putative active promoters as H3K4me3 peaks at transcriptional start sites (TSS) that colocalize with H3K27ac peaks, and putative active enhancers as H3K27ac peaks that were outside of these active promoters. The number of putative enhancers associated with each brain cell type was substantially larger than promoters (microglia: 12,709 and 44,090; neurons: 16,059 and 141,591; oligodendrocytes: 14,370 and 66,525; astrocytes: 14,455 and 63,811 promoters and enhancers, respectively), consistent with a one-to-many relationship between promoters and enhancers. Whereas active promoters are largely shared between cell types (Fig. 1D), a relatively small fraction of active enhancers overlap between cell types (Fig. 1E), indicating that cell type specificity is mainly captured within the enhancer repertoire. Most bulk brain enhancer regions that were identified by PsychENCODE overlapped with the H3K27ac enhancers from at least one of the four cell types (94%). However, more than 87% of cell type-enhancer regions identified in sorted nuclei were not detected by PsychENCODE.

### Gene regulatory regions associated with disease

To determine the enrichment of genetic variants associated with complex traits and diseases in cell type-specific regulatory regions, we performed a linkage disequilibrium score (LDSC) regression analysis of heritability (*27*). LDSC determines whether genetic heritability for a complex trait or disease is enriched for SNPs that fall within specific genome annotations by utilizing GWAS summary statistics and taking into account linkage disequilibrium. We obtained GWAS summary statistics for neurological disorders [AD (*28, 29*), amyotrophic lateral sclerosis (ALS) (*30*), multiple sclerosis (MS) (*31*), Parkinson’s disease (PD)(*32*), and epilepsy (*33*)], psychiatric disorders [autism (*34*), major depressive disorder (MDD) (*35*), bipolar disorder (BD) (*36*), schizophrenia (SCZ) (*37*), and attention deficit hyperactivity disorder (ADHD) (*38*)] and neurobehavior traits [intelligence (*39*), cognitive function (*40*), risk behavior (*41*), neuroticism (*42*) and insomnia (*43*)] (Table S4). LDSC regression analysis was performed for each of these traits and disorders to test whether SNP heritability was enriched in active promoters and enhancers for each cell type, as well as PsychENCODE bulk brain-derived enhancers. Psychiatric disorders and behavioral traits are often assumed to have a neuronal component and, as expected, we found a strong enrichment of heritability for variants within neuronal enhancers for all of these disorders and traits (Fig. 1F). PsychENCODE enhancers had a similar, although reduced, level of heritability for variants within enhancers for psychiatric disorders and behavioral traits (Fig. 1F) (*25*). LDSC analysis of psychiatric disorders and behavioral traits at neuronal promoters showed a comparable enrichment of heritability as variants at neuronal enhancers (Fig. 1F), suggesting that a sizable proportion of GWAS association was possibly driven by dysregulated promoter activity. Furthermore, we found an enrichment of SCZ, BD and behavior trait heritability for genetic variants within oligodendrocyte and astrocyte promoters, suggesting a possible shared role for glia in psychiatric conditions (Fig. 1F). In contrast, AD SNP heritability was most highly enriched in microglia regulatory elements (Fig. 1F). Enrichment of AD SNP heritability was higher in microglia enhancers than promoters and was verified using two independent GWAS studies (*28, 29*), suggesting a prominent role of microglia enhancers in AD pathology (Fig. 1F).

### Inference of cell type-specific, lineage-determining transcription factors

To gain insights into mechanisms driving distinct enhancer landscapes, we performed de novo motif analyses at open chromatin regions associated with distal H3K27ac signal. These analyses identified transcription factor recognition motifs associated with each brain cell type (fig. S3). In addition, the intersection of H3K27ac-defined active promoters with a compilation of human transcription factors found 288 transcription factors that were active in a cell type-specific manner in the brain (fig. S4, Table S3) (*44*). A subset of these cell type-specific transcription factors was paired with enhancer motifs of the corresponding cell type; several of these transcription factors have been associated with disease (Fig. 2, highlighted with an asterisk; Table S5). These criteria confirmed key microglia transcription factors identified by previous epigenetic analyses of ex vivo human microglia, including *SPI1* (encoding PU.1), *IRF8, CEBPA, RUNX1, RUNX2, MEF2C* and *FLI1* (*15*). We extended these analyses to neuronal nuclei and identified motifs for transcription factors involved in neuronal cell fate and cortical layer specification, such as *NEUROD, ASCL* and *TBR1* (*45–47*). In addition, we identified *TWIST2* as a putative neuronal transcription factor; *TWIST2* has been associated with GWAS hits for intellectual disability, hearing impairment, and developmental delay (*44*). We found motifs for factors important for oligodendrocyte specification, such as *SOX6, SOX8 and SOX10* (*48–51*). *SOX10* has been associated with multiple brain conditions, including demyelination disorders (*52*), and *NFIX*, a factor unexplored in oligodendrocytes, has been linked to agenesis of the corpus callosum (*53*). Lastly, we found motifs for transcription factors potentially important for astrocyte identity, such as *SOX9* (*54, 55*).

**Fig. 2.**
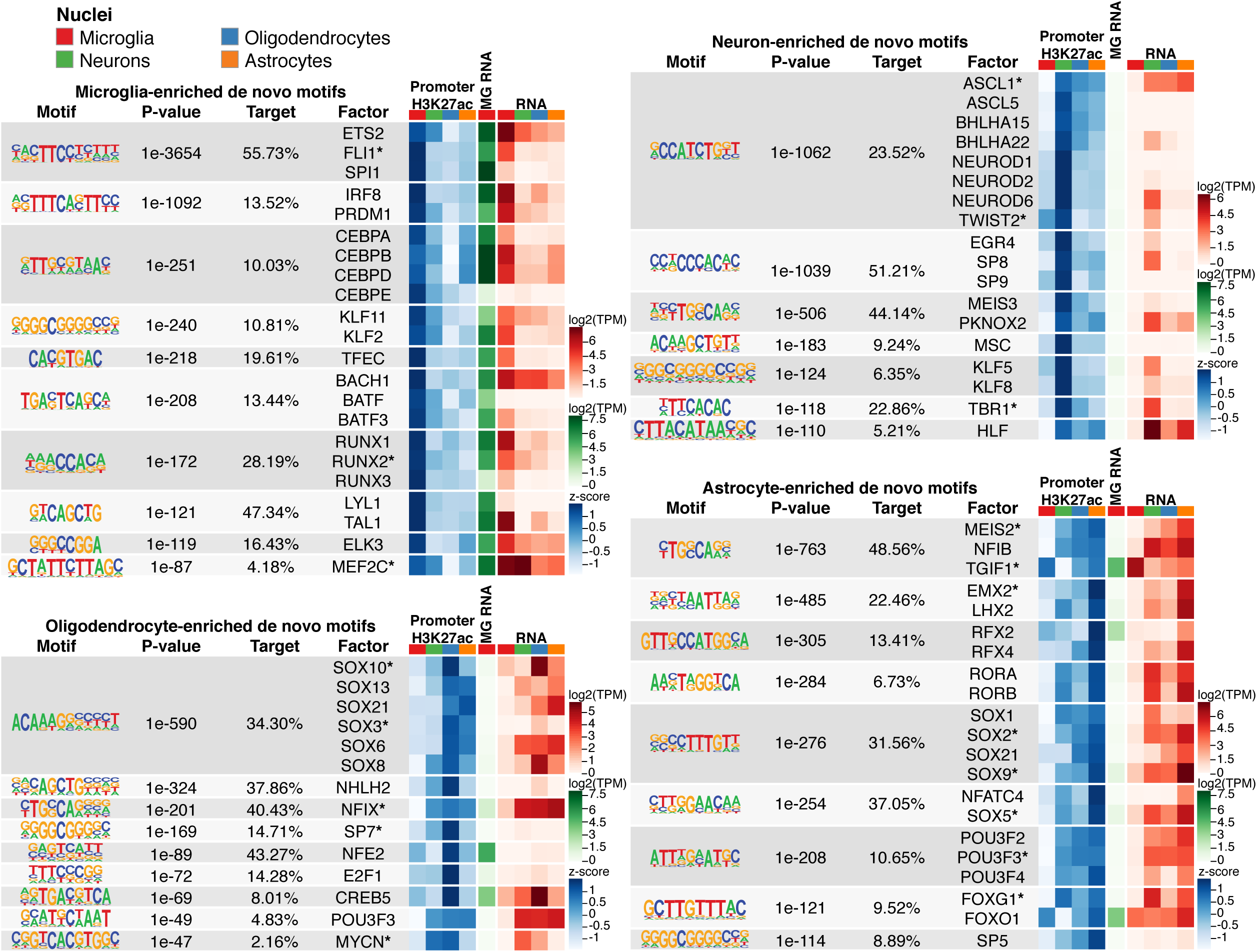
Motif enrichment at active genomic regulatory regions of brain cell types and corresponding candidate transcription factors. Tables show de novo motifs enriched at ATAC-seq peaks associated with H3K27AC-marked enhancers for the indicated brain cell types. Z-score normalized promoter H3K27ac tag counts, ex vivo microglia RNA expression log_2_(transcripts per million (TPM)) (*15*) and brain cell type RNA expression log_2_(TPM) (*21*) for transcription factors predicted to bind the corresponding motifs are shown as heat maps to the right. Transcription factors with an asterisk are associated with GWAS variants identified for brain disorders (Table S5).

### Cell type-specific, promoter-enhancer connectomes

The relationship between promoters and distal regulatory regions for different cell types in the brain is largely unknown. We utilized proximity ligation-assisted ChIP-seq (PLAC-seq) (*56*), a protein-centric chromatin conformation method similar to HiChIP (*57*), in which proximity ligation preceded an enrichment for active promoters by H3K4me3 ChIP-seq. MAPS was used to call chromatin loops between H3K4me3 peaks and distal regulatory regions from PLAC-seq datasets in microglia, neurons and oligodendrocytes (*58*). A representative example is the *SALL1* locus, which has PLAC-seq interactions to cell type-specific enhancers (Fig. 3A). The *SALL1* locus includes a microglia-specific super-enhancer that is PLAC-linked to *SALL1* only in microglia and not the other cell types (Fig. 3A; super-enhancer: highlighted yellow). There were 149,639 significant unique interactions across brain cell types (104,802 microglia; 62,440 neurons; 30,583 oligodendrocytes) (Fig. 3B), which clustered according to cell type of origin (fig. S5A). Most PLAC interactions were classified as chromatin loops that link H3K4me3 peaks to gene regulatory regions (Fig. 3B; microglia: 58.6%; neurons: 62.5%; oligodendrocytes 69.5%; Table S6). A strong H3K4me3 signal in itself does not dictate that an interaction will occur (fig. S5B), suggesting that PLAC-seq captures a unique dimension of the chromatin conformation.

**Fig. 3.**
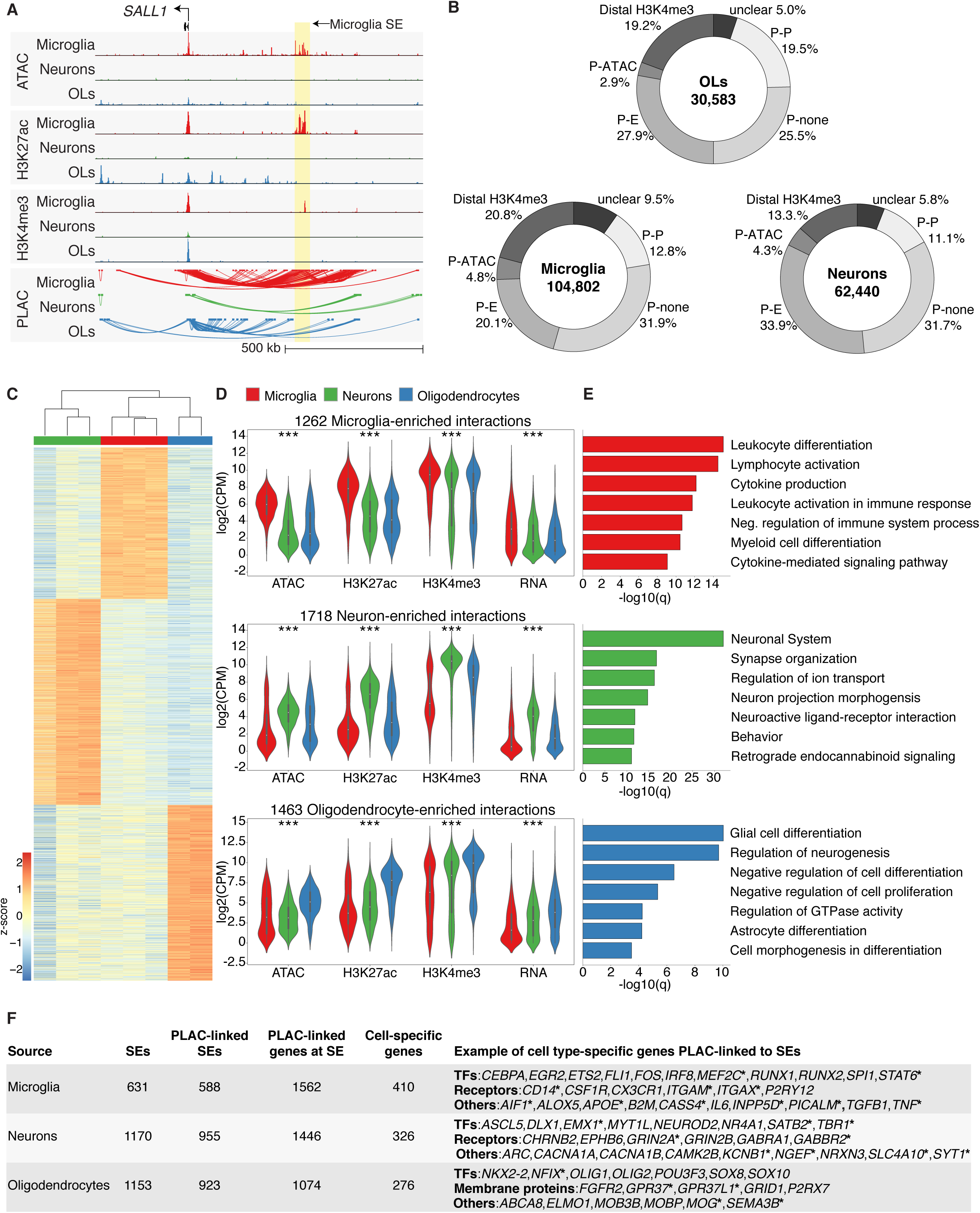
PLAC-seq-defined chromatin loops link promoters to active gene regulatory regions. (A) UCSC browser visualization of open chromatin by ATAC-seq, H3K27ac and H3K4me3 ChIP-seq abundance at gene regulatory regions and promoters, respectively (middle), and PLAC-seq identified chromatin loops (bottom) at the *SALL1* locus. The microglia-specific super-enhancer is highlighted in yellow and is associated with microglia PLAC-seq interactions. (B) Distribution of PLAC-seq interactions at regulatory regions at promoters (‘P’), enhancers (‘E’), open chromatin outside of enhancers (ATAC-seq) and at gene regulatory regions not associated with promoters (distal H3K4me3). (C) Heatmap showing z-score normalized log_2_(counts per million (CPM)) values of PLAC interactions upregulated in microglia, neurons and oligodendrocytes. (D) Violin plots of ATAC-seq, H3K27ac, H3K4me3 ChIP-seq and RNA-seq log_2_(CPM) values at PLAC-seq interactions identified in (C) for microglia, neurons and oligodendrocytes; *** = P < 1e-12; ** = P < 1e-5; * = P < 1e-3. Kruskal-Wallis-between group test. (E) Metascape enrichment analyses of active genes identified at PLAC-seq upregulated interactions shown in (C) for microglia, neurons and oligodendrocytes shown as –log_10_(q) values. (F) Table of super-enhancers linked to cell type-specific genes by PLAC-seq interactions. Asterisks denotes genes with PLAC-seq interactions to super-enhancers that colocalize with disease-risk variants identified by GWAS shown in Fig. 1F (Table S7); SE, super-enhancer.

An analysis of PLAC-seq interactions that were consistently increased between replicates of one cell type compared to the others identified a subset of chromatin connections detected in a cell type-specific manner, either due to cell type-specific chromatin organization or cell type-specific H3K4me3 occupancy of promoters (microglia: 1262, neurons: 1718, oligodendrocytes: 1463 interactions; Fig. 3C). These subsets of PLAC-seq interactions were found to colocalize with increased ATAC-seq, H3K27ac and H3K4me3 ChIP-seq signal in a cell type-specific manner (Fig. 3D). Active promoters linked to microglia, neuronal and oligodendrocyte enriched interactions were associated with gene ontology terms representative of myeloid cells, the nervous system and glia, respectively (Fig 3E). These results illustrate the strength of the PLAC interactome to annotate promoter-enhancer connections in brain cell types.

Super-enhancers are clusters of multiple enhancers that are thought to drive the expression of genes important for cell identity and function (*59*). Using H3K27ac, we identified a total of 2,954 super-enhancers in the microglia, neuron and oligodendrocyte populations, and 83% of these super-enhancers had PLAC interactions in the same cell type (Fig. 3F). These super-enhancers were linked to 4,082 genes of which 1,012 were expressed in a brain cell type-specific manner. Examples of cell type-specific super-enhancer-linked genes that are important for development and function of microglia, neurons and oligodendrocytes are shown in Figure 3F. Many super-enhancers harbor GWAS disease-risk variants and were linked to cell type-specific genes, suggesting that a subset of GWAS variants acts on super-enhancers to affect gene expression (Fig. 3F, denoted with asterisks; Table S7).

### Expansion of the AD gene network

We next used promoter-anchored PLAC-seq data in each cell type to identify connections between promoters and non-coding AD risk alleles (*29*). This integrative approach identified 92 genes that were potentially influenced by these non-coding variants (Table 1), which we categized according to three features. First, we found cell type-specific enhancers harboring AD-risk variants that were PLAC-linked to genes known to be expressed in multiple cell types, implicating a cell type-specific contribution to disease susceptibility. One example is *PICALM*, which, despite being expressed in multiple cell types (*21*), has microglia-specific enhancers harboring AD-risk variants (Fig. 4A), including the rs10792832 lead AD-risk variant for this locus (*29*). Second, we identified AD-risk variants that were PLAC-linked to more distal active promoters and not the closest gene promoter. This has allowed an assignment of distal genes that are more likely to be affected by the GWAS variants than the closest gene. An example is the *SLC24A4* locus, which has a promoter with low activity in all four brain cell types examined but harbors AD-risk variants that were connected to the proximal active promoter of the spinocerebellar ataxia-3 gene, *ATXN3*, as well as *TRIP11* and *CPSF2* (Fig. 4B). Lastly, we observed multiple loci with gene-regulatory regions harboring AD risk variants that were PLAC-linked to active promoters of both GWAS-assigned genes and an extended subset of previously unassigned genes. An example is the *CLU* locus, which has PLAC-linked AD-risk variants to the GWAS-assigned genes *CLU* and *PTK2B*, and an extended set of genes that includes *TRIM35*, *CHRNA2*, *SCARA3* and *CCDC25* (Fig. 4C). Another example is the *KAT8* locus, which has PLAC-linked connections in microglia to the integrins *ITGAX* and *ITGAM* (Table 1); the former has also been implicated by a long-range expression quantitative trait locus (eQTL) association in the peripheral blood with the rs59735493 AD-risk variant (*29*).

**Table 1.**
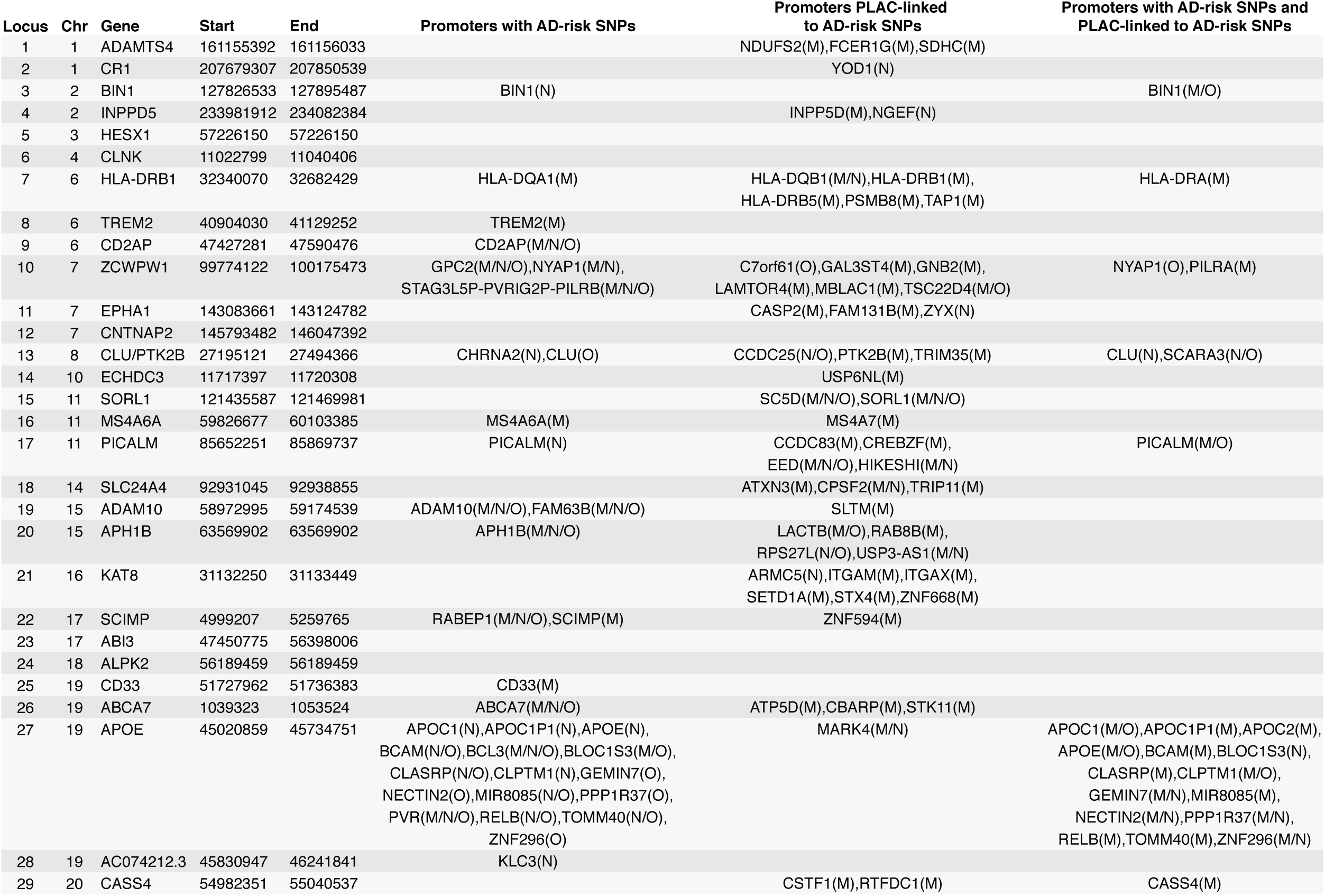
GWAS-identified regions and extended PLAC-seq linked genes assigned to AD-risk loci. Table of genes that have either (1) promoters with AD-risk variants, (2) promoters PLAC-linked to AD-risk variants or (3) promoters with AD-risk variants and PLAC-linked to AD-risk variants. The cell type(s) with active promoters for each gene is indicated for microglia (M), neurons (N) and oligodendrocytes (O). AD SNP information taken from Jansen et al 2019 (29).

**Fig. 4.**
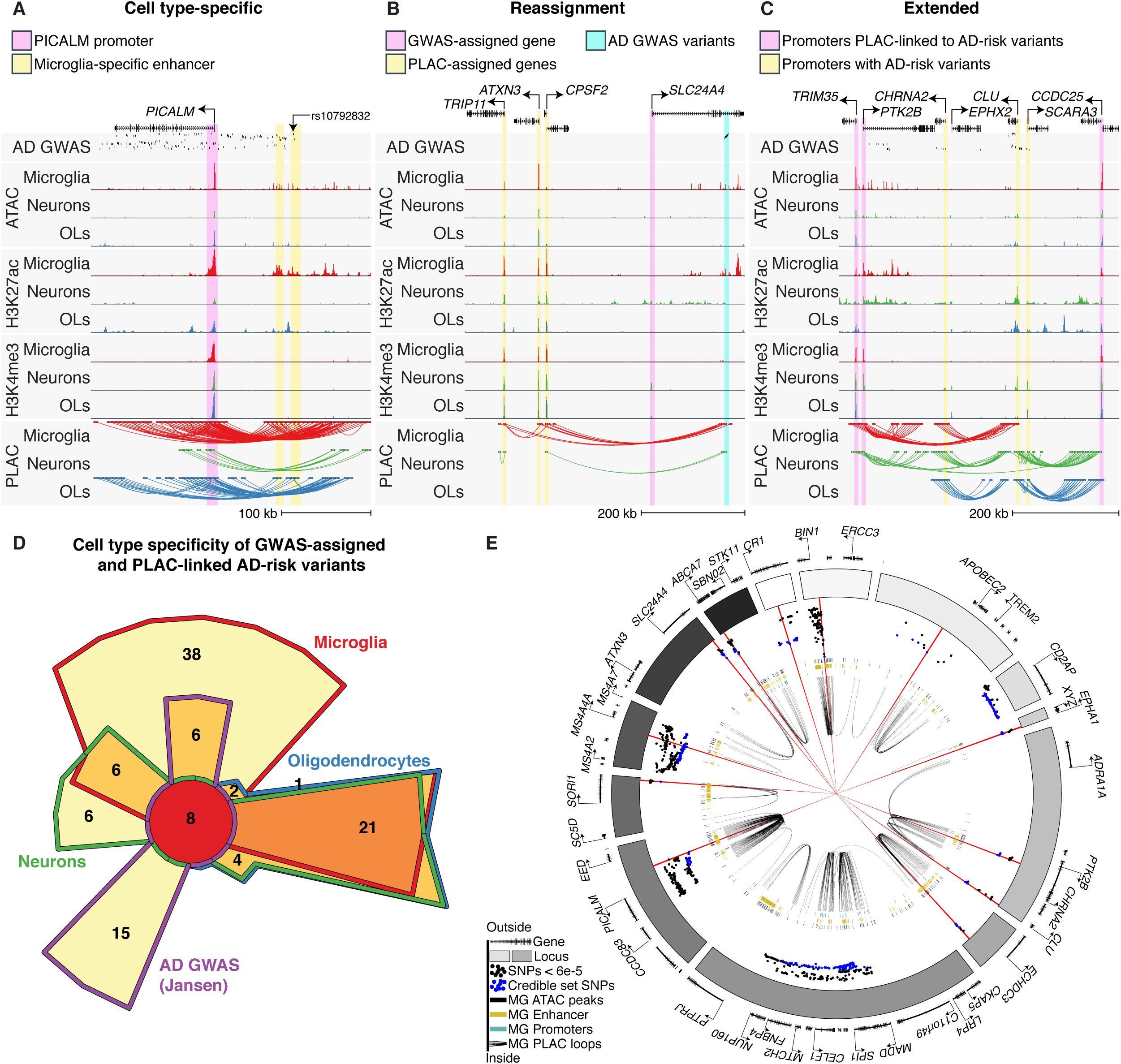
The connectomes of brain cell types at AD-risk loci reveals an expanded gene network. (A)-(C) UCSC genome browser examples of PLAC-seq interactions at AD-risk loci demonstrating (A) cell type-specific gene regulatory regions, (B) reassignment of GWAS-assigned genes and (C) extension of GWAS-assigned genes. (D) Chow-Ruskey plot of GWAS-assigned and PLAC-linked AD-risk variants in microglia, neurons and oligodendrocytes. (E) Circos plot showing genome-wide significant AD loci, microglia enhancer and promoter regions, microglia PLAC-seq loops and high-confidence AD variants identified by fine mapping. Dots represent z-score values of AD SNPs (Kunkle, stage 1) with log10 p-value < 6e-5 (*28*). Blue dots represent z-score values of AD SNPs that were part of the shared credible set SNPs (95% confidence); red lines show 15 high-confidence SNPs with a posterior probability > 0.2. Black bars represent microglia ATAC peaks, gold bars represent microglia enhancers and turquoise bars represent microglia promoters. Black loops represent microglia PLAC-seq connections.

PLAC-seq analysis at AD GWAS loci (*29*) identified 78 genes with microglia active promoters that either overlapped AD-risk variants or were PLAC-linked to AD-risk variants (Fig. 4D; Table 1). Of these 78 genes, 38 had PLAC-seq interactions that were detected in microglia and not the other cell types (Fig. 4D; Table 1). Protein-protein interaction (PPI) network analysis showed that many of the AD-risk genes identified by PLAC-seq were highly connected with GWAS-assigned genes and formed a network centered around APOE (fig. S6A, B), a major genetic risk factor for AD (*60*). AD-risk genes with microglia active promoters were associated with immune function, amyloid-beta processing and neurofibrillary tangle assembly, suggesting that these genes converge on similar pathways that might be amenable to therapeutic targeting (fig. S6C). In contrast, the PPI networks for neuron and oligodendrocyte AD-assigned genes were smaller in scope (fig. S6A). A subset of 15 GWAS-assigned nearest genes were not directly implicated by the epigenomic and connectome analysis (Fig. 4D), suggesting that these genes might not be directly impacted by AD variants in the cell types or cell states we explored.

To get a better understanding of AD genetics and the microglia connectome, we distinguished likely causal variants from those in linkage disequilibrium by applying two complementary fine mapping approaches, CAVIAR (*61*) and PAINTOR (*62*), to a recent AD GWAS (*28*). Fine mapping analysis using PAINTOR was further guided by integrating microglia H3K27ac peaks. The resulting credible sets identified 261 variants that were shared between both fine mapping approaches and ranged from 1 variant at the *SORL1* and *BIN1* loci, to 3 variants at the *PICALM* locus, to 64 variants at the *CD2AP* locus (Fig. 4E; blue circles). In many instances, such as in the *BIN1*, *PICALM*, and *SORL1* loci, the putative fine mapped variants overlapped with microglia-specific enhancers that were linked by PLAC-seq to corresponding gene promoters (Fig. 4E).

### An enhancer harboring a lead AD-risk variant selectively regulates *BIN1* expression in microglia

While virtually all active enhancers are characterized by open chromatin and H3K27ac, not all genomic locations exhibiting these features are essential for gene expression in a particular context. To functionally validate a putative enhancer that contains an AD variant, we chose the putative microglia-specific enhancer containing the AD variant rs6733839 located ∼20 kb upstream of the *BIN1* gene. This variant has an attributable risk percentage for AD of 8.1, which is second only to APOE for late onset idiopathic AD. Furthermore, this variant was identified as a high-confidence variant by fine-mapping (Fig. 4E). This variant is of particular interest because *BIN1* is expressed in neurons, microglia, oligodendrocytes and astrocytes (*21, 63*). Consistent with this finding, H3K27ac and ATAC-seq signal were present at the gene promoters of *BIN1* in microglia, neurons, oligodendrocytes and astrocytes (Fig. 5A; pink highlight). However, enhancer analysis identified putative *BIN1* regulatory regions that were present in microglia but absent in other brain cell types (Fig. 6A; yellow highlight). Additionally, this regulatory region contained the fine-mapped AD-risk variant (rs6733839) and was linked to the BIN1 promoter by microglia PLAC interactions (Fig. 5A; yellow highlight).

**Fig. 5.**
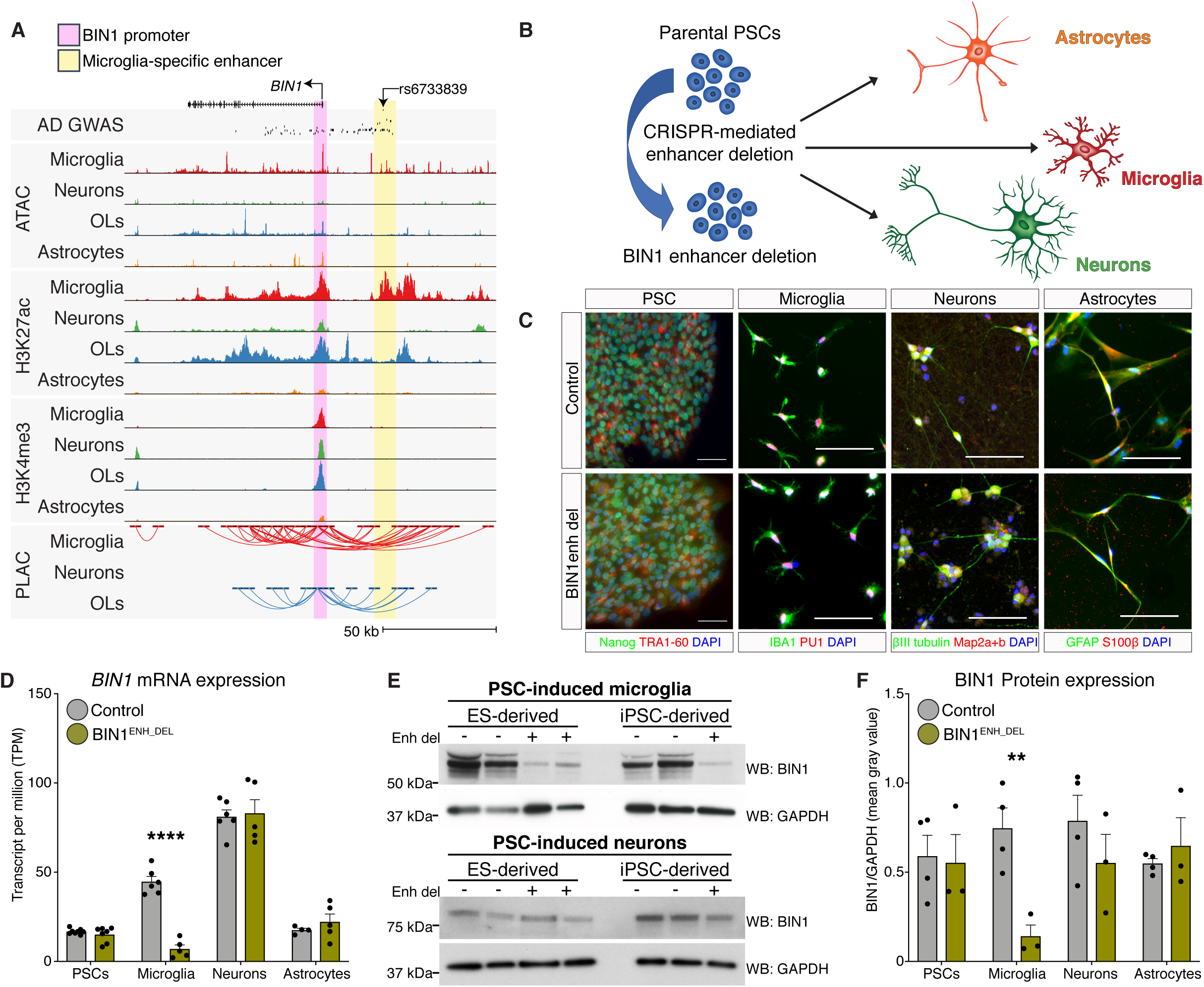
Deletion of a microglia enhancer harboring a lead AD-risk variant affects *BIN1* expression in PSC-derived microglia. (A) UCSC genome browser visualization of AD GWAS risk variants at the *BIN1* locus. Promoter region highlighted in pink shows ATAC-seq, H3K27ac and H3K4me3 ChIP-seq signals in multiple brain cell types. Enhancer region highlighted in yellow shows ATAC-seq and H3K27ac ChIP-seq signal that is restricted to microglia. (B) Schematic of microglia-specific *BIN1* enhancer deletion in PSCs harboring the lead rs6733839 AD-risk variant. Control and enhancer deletion PSCs were differentiated into astrocytes, microglia and neurons. (C) Representative immunohistochemistry images of PSCs, microglia, neurons and astrocytes in control and *BIN1*^enh_del^ lines stained for the indicated cell lineage markers. (D) Gene expression of *BIN1* in control and *BIN1*^enh_del^ PSCs (N = 8,7), microglia (N = 6,5), neurons (N = 6,5) and astrocytes (N = 4,5) as RNA-seq TPM. (E) Western blot of BIN1 and GAPDH in control and *BIN1*^enh_del^ PSC-derived microglia (top) and neurons (bottom). (F) Protein expression of BIN1 in control and *BIN1*^enh_del^ PSCs, microglia, neurons and astrocytes determined as Western blot quantitation of BIN1/GAPDH mean gray intensity. N = 4 controls, 3 *BIN1*^enh_del^ per cell type. **** p-value < 0.0001, ** p-value < 0.01.

Functionality of this microglia-specific, enhancer-promoter interaction was investigated using human pluripotent stem cells (PSCs) to model multiple cell types of the brain in conjunction with CRISPR/Cas9 DNA editing technology (Fig. 5B). We deleted a 363 bp region within the *BIN1* locus using a pair of CRISPR/Cas9 single-guide RNAs in two pluripotent cells lines: an induced pluripotent stem cell (iPSC, EC11) line derived from primary human umbilical vein endothelial cells (*64*) and a human embryonic stem cell line (ESC, H1) (*65*) (fig. S7A). The deleted enhancer region was defined by the borders of a microglia-specific ATAC-seq region previously demonstrated to bind PU.1 (*15*) that includes the rs6733839 AD-risk variant (fig S7B). The normal karyotype of PSC control lines (*BIN1*^control^) and *BIN1* enhancer deletion (*BIN1*^enh_del^) lines were confirmed (fig. S7C). Subsequently, both lines were differentiated into microglia, neurons and astrocytes and showed typical expression of cell lineage markers (Fig. 5C, fig. S8A-D, fig. S9A, B) (*66–68*). Gene expression analysis of *BIN1*^control^ and *BIN1*^enh_del^ lines in PSC and PSC-derived microglia, neurons and astrocytes showed high correlation between samples, with clustering according to cell type (fig. S9C, D). *BIN1* mRNA was expressed in *BIN1*^control^ PSC and PSC-derived cell lines, demonstrating that PSCs are a relevant in vitro model for testing enhancer function of AD-risk genes (Fig. 5D; fig. S9E). Importantly, gene expression of *BIN1* was nearly absent in the *BIN1*^enh_del^ PSC-derived microglia, whereas *BIN1* expression in *BIN1*^enh_del^ PSCs and PSC-derived neurons and astrocytes were equivalent to expression levels in *BIN1*^control^ lines (Fig. 5D; fig. S9E). RNA-seq analysis demonstrated that BIN1 was the only transcript differentially expressed in the *BIN1*^enh_del^ microglia compared to controls, indicating the transcriptional specificity of this microglia enhancer (Table S8). Western blot analysis confirmed BIN1 protein in *BIN1*^control^ PSC-derived microglia cells; however, BIN1 was dramatically reduced in *BIN1*^enh_del^ microglia (Fig. 5E, F). *BIN1* expression was unchanged in the PSC-derived microglia precursor hematopoietic stem cells (HPCs), indicating that the enhancer deletion likely did not affect microglia derivation in vitro (fig S9F). BIN1 protein was detected in *BIN1*^control^ PSCs and PSC-derived neurons and astrocytes and remained unchanged in corresponding *BIN1*^enh_del^ lines (Fig. 5E, F; fig. S9F). These data provide evidence that the genomic region containing rs6733839 functions to selectively regulate BIN1 expression in microglia.

## Discussion

In this study, we established high-quality enhancer and promoter atlases of human microglia, astrocytes, neurons and oligodendrocytes at a resolution beyond that previously generated using bulk tissue. The microglia atlases derived from sorted nuclei were highly similar to those generated from purified microglia sorted from surgically collected brain tissue (*15*), supporting the validity of the approach. Correlation of de novo motif analysis with promoter acetylation enabled inference of transcription factors that are likely to play key roles in the selection and function of enhancers within each cell type. Many of these transcription factors are key regulators of cell identity, such as PU.1 and IRF8 in microglia (*9, 69–74*), ASCL1 and NEUROD1 in neurons (*45–47*), SOX10 and SP7 in oligodendrocytes (*48–50*), and SOX9 and NFIB in astrocytes (*54, 55*). Additionally, we identified many new factors that have not yet been functionally implicated in cell identity, several of which are associated with brain-related disorders. A limitation of the current study is that each cell type exhibits heterogeneity within the brain and the current atlases represent composites that are biased toward their ‘core’ regulatory landscapes. However, the ability to distinguish astrocytes based on a combination of markers suggests that it will be possible to isolate nuclei corresponding to distinct cellular subsets or states for epigenetic analysis.

A recent psychENCODE study, investigating the role of complex genetics in psychiatric diseases, identified almost 80,000 active enhancers from bulk brain tissue of the cortex (*25*). While we were able to recapitulate the majority of psychENCODE enhancers, analysis of cell type-specific nuclei yielded an additional 198,443 cell type specific regulatory regions that were not detectable with bulk tissue. This approach enabled the identification of enrichment for disease heritability to gene regulatory regions for less abundant cell types, such as AD-risk variants in microglia enhancers. We previously found that many genes in the vicinity of AD-risk variants were highly expressed in microglia (*15*), supporting studies that suggested a role for microglia in AD susceptibility (*2, 75–79*). Here we showed that GWAS SNP heritability is enriched in microglia enhancers, providing an association of non-coding genetic variation to AD pathology in microglia. The enrichment of AD risk variants in microglia enhancers does not exclude roles of other cell types for specific AD-risk loci. For example, the *CLU* locus contained active enhancers in neurons, oligodendrocytes and astrocytes but was silent in microglia. Additionally, the complement system has been implicated in microglia-mediated synaptic pruning in AD (*80*). However, the AD locus for the complement receptor 1 (*CR1*) was devoid of active regulatory regions in all the major brain cell types, suggesting that other cell types or cell states also need to be considered.

We also examined heritability of psychiatric disorders and behavioral traits and identified a preferential enrichment in neuronal enhancers. An earlier study comparing neuronal and non-neuronal regulatory regions in the cortex found a similar neuronal enrichment for schizophrenia risk variants (*81*). In addition, we found an enrichment of schizophrenia-risk variants in promoters for neurons, oligodendrocytes and astrocytes, however, whether this reflects contributions of glial to disease or is indicative of a general neural dysfunction remains to be tested.

Chromatin conformation has emerged as a global regulator of cellular function, expression and disease (*10*). Cell type-specific promoter interactome maps thus provide insights into the epigenomic regulation of these cell types by linking putative enhancers and disease variants to target genes. A recent analysis of Hi-C chromatin loops using in vitro iPSC-derived neural progenitor cells, neurons and astrocytes identified dynamic chromatin reorganization during development and was used to generate an extended network of schizophrenia risk genes (*82*). Our approach of using proximity ligation coupled to an enrichment for promoter regions provided an 8.6-fold expansion in the number of detected neuronal chromatin loops. Furthermore, we provide the first description of the promoter interactome in microglia and oligodendrocytes. Using these brain cell type-specific connectivity maps, we provide evidence for microglia-specific targets for a subset of AD GWAS variants, reassigned a subset of variants to targets other than the closest gene, and expanded the assignment of target genes for other variants. While we focused our analysis on genetic variants associated with AD, these datasets can be applied to the complex genetics of psychiatric disorders and neurobehavioral traits. However, future studies of psychiatric disorders will need to examine in vivo developmental regulation of the chromatin interactome.

Combinatorial analysis of enhancer landscapes with promoter interactome maps suggested that the rs6733839 fine-mapped AD-risk variant at the *BIN1* locus resides in a microglia enhancer that is connected to the *BIN1* promoter. *BIN1* has been implicated in diverse cellular processes such as endocytosis, actin dynamics, DNA repair, membrane trafficking, inflammation and apoptosis (*28*), and regulation of tau pathology in disease (*83*). Prior studies suggested that AD variants at the *BIN1* locus played a putative role in neurons (*83, 84*) and oligodendrocytes (*85*). However, we recently showed that BIN1 is highly expressed in ex vivo microglia compared to the whole cortex and has downstream active enhancers bound by the lineage-determining transcription factor PU.1 (*15*). Subsequently, a study found cell type-specific protein expression of *BIN1* isoforms in microglia, as well as in other cell types of the brain (*63*). Here, we showed that the *BIN1* locus has enhancer landscapes unique to each cell type and that the *BIN1*-linked enhancer harboring the rs6733839 AD-risk variant is highly active in microglia compared to other cell types. Deletion of the *BIN1* microglia enhancer greatly reduces *BIN1* expression in PSC-derived microglia without effecting gene expression in neurons or astrocytes. Thus, an AD-risk gene that is expressed in multiple cell types may have a microglia-specific propensity for pathology through localization of disease-risk variants to cell type-specific enhancers. However, the function of the rs6733839 AD-risk variant on enhancer activity and gene transcription remains to be explored in microglia. Our previous studies demonstrated that *BIN1* expression in ex vivo microglia was dependent on the brain environment. It is thus possible that effects of rs6733839 on enhancer activity will be dependent on an in vivo context, perhaps in association with amyloid or tau pathology. Recent studies indicate that iPSC-derived human microglia engrafted into the mouse brain humanized for CSF-1 assume an in vivo phenotype (*86, 87*), suggesting a future approach to establish functional consequences of non-coding variants in human microglia.

The present studies have relied on freshly frozen, surgically collected brain tissue to define an initial set of atlases of the enhancer and promoter landscapes of specific brain cell types and their interconnections. These atlases are likely to provide valuable resources for investigation of diverse questions related to human neurodegenerative and behavioral disorders. In addition, the associated methodologies set the stage for investigation of how these transcriptional regulatory landscapes are altered in a disease-specific manner using the substantial existing repositories that store freshly frozen brain tissue from patients and control subjects. Examination of the cell type-specific changes in the enhancer landscapes of these tissues is likely to lead to new insights into transcription factors and upstream signaling pathways that drive pathogenic programs of gene expression.

## Acknowledgments

We thank J. Collier for technical assistance, L. Van Ael for assistance with manuscript preparation, D. Skola, C. Han and M. L. Gage for assistance with manuscript editing, C. Benner for discussions on motif analysis and M. Gymrek for discussions on fine mapping.

## Funding

These studies were supported by the NIH grants RF1 AG061060-01, R01 AG056511-01A1, R01 AG057706-01, and a Cure Alzheimer’s Fund Gifford Neuroinflammation Consortium. AN was supported by a grant from the Alzheimer’s Association (AARF-18-531498). AN was partially supported by the Altman Clinical & Translational Research Institute (ACTRI) at UCSD. The ACTRI is funded from awards issued by the National Center for Advancing Translational Sciences, NIH KL2TR001444. I.R.H. was supported by the VENI research program, which is financed by the Netherlands Organization for Scientific Research (NWO). NGC was funded by NIH K08 NS109200-01 and by the Hartwell Foundation. C.Z.H. is supported by the Cancer Research Institute Irvington Postdoctoral Fellowship Program. CO was funded by NIH-NCI CCSG: P30 014195 and S10-OD023689. Additional support was provided by the JPB Foundation, The Engman Foundation, The AHA/ Allen Award, and the Dolby Foundation. J.B.B. and R.A.S. were partially supported by AG062429-01.

## Author contributions

A.N., N.G.C., J.C.M.S., and C.K.G. conceived the study. N.G.C. coordinated tissue acquisition, clinical chart review, and institutional review board approval. D.D.G., and M.L.L. obtained patients’ informed consent, and resected and evaluated brain tissue. A.N. and J.C.M.S. optimized isolation protocol. A.N. isolated nuclei, with assistance from M.P.P. with nuclei sorting and the analysis of flow cytometry data. D.G. isolated ex vivo microglia used for PLAC-seq and H3K4me3 ChIP-seq. N.G.C. performed PSC experiments with assistance from M.P., J.X., Y.W., Z.K for cell culture and assistance from C.O for cell sorting. A.N. prepared ChIP-seq and ATAC-seq libraries. A.N. and I.R.H. preprocessed and analyzed ChIP-seq, ATAC-seq and RNA-seq datasets. I.R.H. performed motif analysis with assistance from Z.S for analysis. I.R.H. analyzed PLAC-seq and GWAS datasets. M.Y. and R.H. prepared PLAC-seq libraries, with B.R for supervision. A.N., I.R.H., C.K.G. wrote the manuscript, with contributions from N.G.C., C.Z.H., J.C.M.S., F.H.G. and B.R.

## Competing interests

B.R. is co-founder of Arima Genomics, Inc, which sells Hi-C and PLAC-seq kits.

## Data and Material

Primary data is being prepared for submission to the dbGAP data base for access upon registration.

## Supplemental Materials

### Methods and Materials

#### Human tissue collection

Nuclei were isolated from human brain tissue resected for treatment of epilepsy in 9 individuals (Table S1). Brain tissue was obtained with informed consent under a protocol approved by the UC San Diego and Rady Children’s Hospital Institutional Review Board. The reported data sets are from sequential samples for which sequencing libraries met technical quality standards. Resected brain tissue was immediately placed on ice, transferred to the laboratory, and either snap frozen and stored at -80 °C prior to nuclei isolation, or immediately processed for isolation of ex vivo live microglia. Nuclei were collected to generate 4x PU.1, 4x NEUN, 4x OLIG2 and 3x LHX2 H3K27ac ChIP-seq datasets (Table S1). Nuclei were collected to generate 3x PU.1, 4x NEUN, 3x OLIG2 and 3x LHX2 ATAC-seq datasets (Table S1). Nuclei were collected to generate 1x PU.1, 2x NEUN, 3x OLIG2 and 2x LHX2 H3K4me3 ChIP-seq datasets and ex vivo microglia were harvested to generate 1x microglia H3K4me3 ChIP-seq dataset (Table S1). Nuclei were collected to generate 3x NEUN and 2x OLIG2 PLAC-seq datasets and ex vivo microglia were harvested to generate 3x microglia PLAC-seq datasets (Table S1).

#### Nuclei isolation with FANS

Frozen brain tissue was homogenized in 1% formaldehyde in Dulbecco’s phosphate buffered saline (PBS; Corning) and incubated at room temperature for 10 mins. Fixed tissue was quenched with 0.125 M glycine for 5 mins at room temperature and pelleted at 1,100 xg for 10 mins. Subsequent steps were performed on ice or at 4°C. Homogenates were washed two times with NF1 buffer (10 mM Tris-HCl pH 8.0, 1 mM EDTA, 5 mM MgCl_2_, 0.1 M sucrose, 0.5 % Triton X-100). Homogenates were incubated on ice in 5 ml NF1 for 30 mins. Nuclei were extracted from homogenates using a glass Dounce and passed through a 70-μm strainer. Homogenates were underlaid with a 1.2 M sucrose cushion and centrifuged at 4,000 xg for 30 mins. Pelleted nuclei were washed with NF1 buffer at 1,600 xg for 5 mins. Nuclei were washed with staining buffer (PBS, 1 % bovine serum albumin, 1 mM EDTA) at 1,600 xg for 5 mins. Nuclei were suspended in 0.5 ml staining buffer and incubated overnight at 4°C with the following antibodies: PU.1 Alexa Fluor 647 (1:500; BioLegend, 658004), NEUN Alexa Fluor 488 (1:2,500; Millipore Sigma, MAB377), OLIG2 (1:1,000; Abcam, ab109186) and LHX2 (1:500; Abcam, ab219983). Nuclei were washed with staining buffer. OLIG2 and LHX2 unconjugated antibodies were subsequently stained with goat anti-rabbit Alexa Fluor 647 (1:4,000; ThermoFisher Scientific, A21244) and washed with staining buffer. Nuclei were passed through a 30-μm strainer and stained with 0.5 μg ml^−1^ DAPI. Nuclei for each cell type were sorted with a Beckman Coulter MoFlo® Astrio™ EQ cell sorter and pelleted at 1,600 xg for 15 mins in staining buffer. Nuclei for ChIP-seq and PLAC-seq were stored at -80°C prior to library preparation; nuclei for ATAC-seq were immediately processed.

#### Ex vivo microglia isolation

Microglia isolation was performed as previously described (*15*).

#### Chromatin immunoprecipitation sequencing (ChIP–seq)

Nuclei (500,000 +/- 50,000) were thawed on ice, resuspended in 130 μl LB3 (10 mM Tris-HCl pH 7.5, 100 mM NaCl, 1 mM EDTA, 0.5 mM EGTA, 0.1% Na-Deoxycholate, 0.5% N-Lauroylsarcosine, 1x protease inhibitors) and transferred to microtubes with an AFA Fiber (Covaris, MA). All subsequent steps were performed on ice or at 4°C. Samples were sonicated using a Covaris E220 focused-ultrasonicator (Covaris, MA) for 240 secs (Duty: 5, PIP: 140, Cycles: 200, AMP/Vel/Dwell: 0.0). Samples were adjusted to 250 μl with 1% Triton X-100, centrifuged at 21,000 xg for 10 mins and the pellet was discarded. 1 % of the sample was stored at -20°C for DNA input control. For ChIP, the following were added and rotated at 4°C overnight: 25 μl Protein G Dynabeads and either H3K27ac antibody (2 μl serum; Active Motif, 39135) or H3K4me3 antibody (2 μl; Millipore Sigma, 04-745). Dynabeads were washed 3 times with Wash Buffer 1 (20 mM Tris-HCl pH 7.4, 150 mM NaCl, 2 mM EDTA, 0.1% SDS, 1% Triton X-100), three times with Wash Buffer 3 (10 mM Tris-HCl pH, 250 mM LiCl, 1 mM EDTA, 1% Triton X-100, 0.7% Na-Deoxycholate) and three times with TET (10 mM Tris-HCl pH 8, 1 mM EDTA, 0.2% Tween20). Dynabeads were washed once with TE-NaCl (10 mM Tris-HCl pH 8, 1 mM EDTA, 50 mM NaCl) in PCR tubes and resuspended in 25 μl TT (10 mM Tris-HCl pH 8, 0.05% Tween20). Input samples were adjusted to 25 μl with TT and libraries were generated in parallel with ChIP samples. Library NEBNext End Prep and Adaptor Ligation were performed using NEBNext Ultra II DNA Library Prep kit (New England BioLabs) according to manufacturer instructions using barcoded adapters (NextFlex, Bioo Scientific). Libraries were PCR amplified for 14 cycles with NEBNext High Fidelity 2X PCR MasterMix (New England BioLabs, NEBM0541). Libraries were size selected for 225 – 350 bp fragments by gel extraction (10% TBE gels, Life Technologies) and were single-end sequenced for 51 cycles on an Illumina HiSeq 4000 (Illumina, San Diego, CA).

#### Assay for Transposase-Accessible Chromatin-sequencing (ATAC-seq)

We performed ATAC-seq on fixed nuclei samples using a published protocol with slight modifications (*22*). Immediately post-sorting, nuclei (200,000 +/- 20,000) were pelleted at 1,600 xg for 15 mins at 4°C. Pellets were resuspended in lysis buffer (10 mM Tris-HCl pH 7.4, 10 mM NaCl, 3 mM MgCl_2_, 0.5% IGEPAL CA-630) and centrifuged at 1,600 xg for 10 mins. Pellets were resuspended in 50 μl transposition mix (1x Tagmentation buffer (Illumina, 15027866), 2.5 μI Tagmentation DNA Enzyme I (Illumina, 15027865) in 50 μI reaction) and incubated at 37°C for 30 mins. Samples were incubated with 150 μl reverse-crosslinking solution (50 mM Tris-HCl pH 7.4, 200 mM NaCl, 1 mM EDTA, 1% SDS) at 65°C overnight. Samples were incubated with 1 μl Proteinase K (5 ng ml^−1^) at 55°C for 2 hours. DNA was purified using ChIP DNA and Clean Concentrator kit (Zymo Research; D5205). Libraries were PCR amplified for 8-12 cycles with NEBNext High Fidelity 2X PCR MasterMix (New England BioLabs, NEBM0541) and 1.25 μM Nextera index primer 1 and 1.25 μM Nextera index primer 2-barcode. Libraries were size selected for 155 – 250 bp fragments by gel extraction (10% TBE gels, Life Technologies) and were single-end sequenced for 51 cycles on an Illumina HiSeq 4000 (Illumina, San Diego, CA).

#### Ribonucleic acid-sequencing (RNA-seq)

PSCs and PSC-derived cells (100,000) were collected in 600 μl Trizol and RNA was purified with Zymo kit (Zymo Research) and eluted in 50 μl water. mRNA was isolated by incubating with 10 μl oligo(dT) beads in 50 μl 2x DTBB (20 mM Tris-HCl pH 7.5, 1 M LiCl, 2 mM EDTA, 1 % lithium dodecyl sulfate, 0.1 % triton X-100) at 65°C for 2 mins, followed by an incubation at room temperature for 10 mins, 1 wash with Wash Buffer 1 (10 mM Tris-HCl pH 7.5, 0.15 M LiCl, 1 mM EDTA, 0.1 % lithium dodecyl sulfate, 0.1 % triton X-100) and 1 wash with Wash Buffer 3 (10mM Tris-HCl pH 7.5, 0.15 M NaCl, 1 mM EDTA). mRNA was eluted in 50 μl elution buffer (10mM Tris-HCl pH 7.5, 1mM EDTA) at 80°C for 2 mins. mRNA isolation was repeated using fresh oligo(dT) beads and was subjected to a final wash with 1x SuperScript III first-strand buffer. mRNA was eluted from oligo(dT) beads and fragmented in 10 μl 2x SuperScript III at 94°C for 9 mins. Isolated mRNA was used for first-strand cDNA synthesis as follows: 1.5 μg random primers (ThermoFisher Scientific, 48190011), 2 μM oligo d(T) (ThermoFisher Scientific,18418020), 10 U SUPERase-In (AM2696) and 0.8 mM dNTP were added to final volume of 12.5 μl; samples were incubated at 50°C for 1 min; 0.2 μg Actinomycin D (Millipore Sigma, A1410), 5 mM DTT and 0.01% Tween20 and 100 U SuperScript III (ThermoFisher Scientific, 18080044) were added to final volume of 20 μl; samples were incubated at 25°C for 10 mins and 50°C for 50 mins and then on ice. Samples were purified using RNAClean XP beads (Beckman Coulter, A63987) and eluted with 10 μl water. Second-strand synthesis was performed as follows: 1X Blue buffer (Enzymatics, B0110), 0.66 mM PCR mix (Affymetrix, 77330), 0.66 mM dUTP (Affymatrix, 77206), 1 U RNase H (Enzymatics, Y9220L), 10 U DNA polymerase I (Enzymatics, P7050L) were added to a 15 μl final volume; samples were incubated at 16°C for 2.5 hours. cDNA was purified using 2 μl SpeedBeads (ThermoFisher Scientific, 651520505025) in 28 μl 20% PEG8000/2.5 M NaCl and eluted with 40 μl EB (Zymo Research). Samples were subjected to dsDNA repair using 1x T4 ligase buffer, 0.2 mM dNTP, 0.9 U T4 DNA polymerase (Enzymatics, P7080L), 0.3 U Klenow Fragment (Enzymatics, P7060L), 3 U T4 PNK (Enzymatics, Y9040L) in a final volume of 50 μl and incubated at 20°C for 30 mins. Samples were purified using 2 μl SpeedBeads in 93 μl 20% PEG8000/2.5 M NaCl and eluted with 15 μl EB. Samples were subjected to dA-tailing using 1x Blue buffer (Enzymatics, B0110), 0.1 mM dATP, 1.5 U Klenow 3’-5’ exo- (Enzymatics, P7010-LC-L) in a final volume of 30 μl and incubated at 37°C for 30 mins. Samples were purified using 2 μl SpeedBeads in 55.8 μl 20% PEG8000/2.5 M NaCl and eluted with 14 μl EB. Samples were subjected to Y-shape adapter ligation using 0.8 μM barcoded adapters (NextFlex, Bioo Scientific), 1x Rapid Ligation buffer (Enzymatics, B1010L), 300 U T4 DNA Ligase HC (Enzymatics, L6030-HC-L) in a final volume of 30 μl and incubated at 21°C for 15 mins. Samples were purified using 2 μl SpeedBeads in 7.5 μl 20% PEG8000/2.5 M NaCl and eluted with 14 μl EB. Samples were second strand digested with 2 U UDG (Enzymatics, G5010L) at 37°C for 30 mins. Libraries were PCR amplified for 12-16 cycles with the KAPA HiFi HotStart PCR kit (ThermoFisher Scientific, KK2502). Libraries were size selected for 200 – 475 bp fragments by gel extraction (10% TBE gels, Life Technologies) and were single-end sequenced for 51 cycles on an Illumina HiSeq 4000 (Illumina, San Diego, CA).

#### Proximity Ligation ChIP-sequencing (PLAC-seq)

PLAC-seq libraries were prepared for ex vivo microglia, NEUN nuclei and OLIG2 nuclei as previously described with minor modifications (*56, 88*). In brief, purified populations were cross-linked for 15 minutes at room temperature with 1% formaldehyde and quenched for 5 mins at room temperature with 0.2 M glycine (Thermo Fisher Scientific). The cross-linked cells/nuclei were centrifuged at 2,500 xg for 5 mins. To isolate nuclei, cross-linked cells were resuspended in 200 μL lysis buffer (10 mM Tris-HCl (pH 8.0), 10 mM NaCl, 0.2% IPEGAL CA-630) and incubated on ice for 15 minutes. The suspension was then centrifuged at 2,500 xg for 5 mins and the pellet was washed by resuspending in 300 μL lysis buffer and centrifuging at 2,500 xg for 5 mins. The pellet was resuspended in 50 μL 0.5% SDS and incubated for 10 mins at 62°C. 160 μL 1.56% Triton X-100 was added to the suspension and incubated for 15 mins at 37°C. 25 μl of 10X NEBuffer 2 and 100 U MboI were added to digest chromatin for 2 hours at 37°C with rotation (1,000 rpm). Enzymes were inactivated by heating for 20 mins at 62°C. Fragmented ends were biotin labeled by adding 50 μL of a mix containing 0.3 mM biotin-14-dATP, 0.3 mM dCTP, 0.3 mM dTTP, 0.3 mM dGTP, and 0.8 U μl^−1^ Klenow and incubated for 60 mins at 37°C with rotation (900 rpm). Ends were subsequently ligated by adding a 900 μL master mix containing 120 μL 10X T4 DNA ligase buffer (NEB), 100 μL 10% TritionX-100, 6 μL 20mg ml^−1^ BSA, 10 μL 400 U μl^−1^ T4 DNA Ligase (NEB, high concentration formula) and 664 μL H_2_O and incubated for 120 mins at 23°C with 300 rpm slow rotation. Nuclei were pelleted for 5 mins at 4°C with centrifugation at 2,500 xg. For the ChIP, nuclei were resuspended in RIPA Buffer (10 mM Tris (pH 8.0), 140 mM NaCl, 1 mM EDTA, 1% Triton X-100, 0.1% SDS, 0.1% sodium deoxycholate) with proteinase inhibitors and incubated on ice for 10 mins. Sonication was performed using a Covaris M220 instrument (Power 75W, duty factor 10%, cycle per bust 200, time 10 mins, temperature 7°C) and nuclei were spun for 15 mins at 14,000 rpm at 4°C. 5% of supernatant was taken as input DNA. To the remaining cell lysate was added anti-H3K4me3 antibody-coated Dynabeads M-280 Sheep anti-Rabbit IgG (5 μg antibody per sample, Millipore, 04-745), followed by rotation at 4°C overnight for immunoprecipitation. The sample was placed on a magnetic stand for 1 min and the beads were washed three times with RIPA buffer, two times with high-salt RIPA buffer (10 mM Tris pH 8.0, 300 mM NaCl, 1 mM EDTA, 1% Triton X-100, 0.1% SDS, 0.1% deoxycholate), one time with LiCl buffer (10 mM Tris (pH 8.0), 250 mM LiCl, 1 mM EDTA, 0.5% IGEPAL CA-630, 0.1% sodium deoxycholate) and two times with TE buffer (10 mM Tris (pH 8.0), 0.1 mM EDTA). Washed beads were treated with 10 μg RNase A in extraction buffer (10 mM Tris (pH 8.0), 350 mM NaCl, 0.1 mM EDTA, 1% SDS) for 1 hour at 37°C, and subsequently 20 μg proteinase K was added at 65°C for 2 hours. ChIP DNA was purified with Zymo DNA clean & concentrator-5. For Biotin pull down, 25 μL of 10 mg ml^−1^ Dynabeads My One T1 Streptavidin beads was washed with 400 μl of 1X Tween Wash Buffer (5 mM Tris-HCl (pH 7.5), 0.5 mM EDTA, 1 M NaCl, 0.05% Tween) supernatant removed after separation on a magnet. Beads were resuspended with 2X Binding Buffer (10 mM Tris-HCl (pH 7.5), 1 mM EDTA, 2 M NaCl), added to the sample and incubated for 15 mins at room temperature. Beads were subsequently washed twice with 1X Tween Wash Buffer and in between heated on a thermomixer for 2 mins at 55°C with mixing and washed once with 1X NEB T4 DNA ligase buffer. Library prep was prepared using QIAseq Ultralow Input Library Kit (Qiagen, 180492). KAPA qPCR assay was performed to estimate concentration and cycle number for final PCR. Final PCR was directly amplified off the T1 beads according to the qPCR results, and DNA was size selected with 0.5X and 1X SPRI Cleanup and eluted in 1X Tris Buffer and paired-end sequenced.

#### Human pluripotent stem cell culture

All studies were conducted according to the human stem cell (hESCRO) protocol approved by the Salk Institute for Biological Studies. Human embryonic stem cell (ESC) line H1 (WiCell Research Institute, Madison, WI) (*65*) and induced pluripotent stem cell (iPSC) line EC11, derived from primary human umbilical vein endothelial cells (Lonza, Bioscience) (*64*), were cultured utilizing standard techniques. In brief, cells were cultured in StemMacs iPS-Brew media (Miltenyi Biotech, Auburn, CA) and routinely passaged utilizing Gentle Cell Dissociation Reagent (STEMCELL Technologies) onto Matrigel-coated (1 mg ml^−1^) plates. Karyotype was established by standard commercial karyotyping (WiCell Research Institute, Madison, WI).

#### CRISPR/Cas9 gene editing

CRISPR guides were designed utilizing ChopChop v2 (*89*). BIN1 enhancer guides: GTAAGTCACTGGCTATGCATAGG and UAGAGUGCCUGCUCAAACGCAGG. ESC/iPSC CRISPR was performed as follows. Guide RNAs (sgRNA) were prepared by dissolving in nuclease-free TE buffer (IDT Technologies) at a final concentration of 100 μM, then complexed with Cas9 (NEB) at a ratio of 2:3 for 10 mins at room temperature. Pluripotent cells were harvested as single cells utilizing Tryple (Gibco). Cells were counted with live/dead discrimination using trypan blue (Sigma), and 150,000 cells were utilized per electroporation. Cells were electroporated utilizing an Amaxa electroporator (Lonza Biosciences) and the Human Stem Cells Nucleofector 2 Kit (Lonza). Each electroporation was plated into 1 well of a 6-well Matrigel-coated plate with 10 μM ROCK inhibitor. After recovery, cells were plated at a low cell concentration as single cells using Tryple for single colony isolation. Once single colonies were obtained, cells were incubated in collagenase IV (Invitrogen) until colonies were free-floating, and individual colonies were plated to a 96-well Matrigel coated plated with 10 μM ROCK inhibitor. Colonies were then screened for the enhancer deletion. Controls were isolated from the same sub-cloning experiment, which did not have an enhancer deletion. CRISPR deletion was confirmed utilizing the following primers outside of the enhancer deletion region. Fwd: CACGGGTGCAGATGGAAT and rev: AGGGCTGGTGTGAAGGTTA, which yielded a 605 bp product if unedited and approximately 242 bp product if the enhancer was deleted. Enhancer deletions were confirmed by isolating the edited product band and cloning into TOPO-TA (Invitrogen) and sequencing the resulting plasmid to confirm deletion. All of the individually isolated deletion lines: ESC (2) and iPSC (1) had the same 363bp deletion: CCTATGCATAGCCAGTGACTTACGCTGACTTTTCTGGAAACTGGGACTTTAAATAGGAGTTTTTAAAAAGGAAAGAAATCTCTGTTCTGCTTCTTAAAAACACCCTTTTCCCCTTTTTACTTTCAGAAGAAGGAACCAGAGAAGACTAGAAACAGGATGGAACGGGAAGGTGGAGGGAGAGGCGAAGGGGAAGTCTCGATGCTGACTACAAGGCCCCCTCATCCTCCGGCCTCCGGCTTGCAGGGACTGCCCCTGCCTTGGGAGAGTCCTCACAGGCCACACCTGCCACTGGGCCAGGCCCAGAGCAGACACTCACGACAGATGATAAGAATGACAGCTGCCTGCGTTTGAGCAGGCACTCTA.

#### Pluripotent stem cell differentiation

Microglia were generated as previously described with minor modifications (*66*). Briefly, ESC/iPSCs were plated in iPS-Brew with 10 μM ROCK inhibitor (Stem Cell Technologies) onto Matrigel-coated (1 mg ml^−1^) 6-well plates (Corning) using ReLeaSR (STEMCELL Technologies). Cells were differentiated to CD43+ hematopoietic progenitors using the StemCell Technologies Hematopoietic Kit (Cat #05310). On day 1, cells were changed to basal media with supplement A (1:200), supplemented with an additional 1 ml/well on day 3, and changed to basal media with supplement B (1:200) on day 3. Cells received an additional 1 ml/well of medium B on days 5, 7, and 10. Nonadherent hematopoietic cells were collected on days 11 and 13. Cells were then replated onto Matrigel-coated plates (1 mg ml^−1^) at a density of 300,000 cells/well in microglia media. Microglia media consisted of DMEM/F12 (Thermofisher), 2x insulin-transferrin-selenite (Gibco), 2x B27 (Lifetech), 0.5x N2 (Lifetech), 1x Glutamax (Gibco), 2x non-essential amino acids (Gibco), 400 μM monothioglycerol, and 5 μg ml^−1^ insulin (Sigma). Microglia media was supplemented with 100 ng ml^−1^ IL-34 (Proteintech), 50 ng ml^−1^ TGFβ1 (Proteintech) and 25 ng ml^−1^ M-CSF (Proteintech). Cells were supplemented with microglia media with IL-34, TGFβ1 and M-CSF, with removal of media (1 ml remaining) halfway through differentiation. 25 days after initiation with microglia media, cells were resuspended in microglia media with IL-34, MCSF and TGFβ1 with the addition of CD200 100 ng ml^−1^ (Novoprotein) and CX3CL1 100 ng ml^−1^ (Peprotech). Cells were collected on Day 28 for further experiments.

Astrocyte differentiations were performed as previously described through a glial progenitor cell (GPC) intermediate (*67*) with modifications. Briefly, embryoid bodies were prepared from confluent stem cell cultures by mechanical dissociation with 1 mg ml^−1^ collagenase IV (Invitrogen), plated onto ultra-low attached plates in iPS-Brew media supplemented with 10 μM ROCK inhibitor (Stem Cell Technologies) and incubated overnight with agitation. Cells were then transitioned to astrocyte medium (ScienCell) supplemented with 500 ng ml^−1^ Noggin (R&D) and 10 ng ml^−1^ PDGFAA (Peprotech) for a total of 14 days. Thereafter cells were cultured in astrocyte media with only PDGFAA for a further week. At the conclusion, embryoid bodies were dissociated with papain (Worthington) and the resulting GPCs were expanded on 10 μg ml^−1^ poly-L-ornithine-(Sigma) and 1 μg ml^−1^ laminin-coated (Invitrogen) plates in astrocyte media supplemented with 20 ng ml^−1^ FGF2 (Joint Protein Central) and 20 ng ml^−1^ EGF2 (Humanzyme). Astrocyte differentiation was completed utilizing a serum-free system (*21*). Specifically, GPCs were cultured for 4 weeks in Sato media, which consisted of a 1:1 mixture of DMEM (Thermo Fisher) and Neurobasal media (Thermo Fisher) supplemented with 1x PenStrep (ScienCell), 1 mM of sodium pyruvate (Sigma), 2 mM L-glutamine (Life Sciences), 5 μg ml^−1^ N-acetyl-L-cysteine (Sigma), 5 ng ml^−1^ EGF, 20 ng ml^−1^ CNTF (Proteintech), 20 ng ml^−1^ BMP4-4 (Peprotech), and 4 μg ml^−1^ insulin (Sigma). Sato media was further supplemented with 100 μg ml^−1^ of transferrin (Sigma), 100 μg ml^−1^ bovine serum albumin (Sigma), 16 μg ml^−1^ putrescine (Sigma), 0.2 μM progesterone, and 40 ng ml^−1^ sodium selenite (Sigma). Cultures were maintained on the aforementioned laminin and poly-L-lysine-coated plates throughout.

Neurons were differentiated directly from ESCs/iPSCs as previously described (*68*) with modifications. Briefly, ESC/iPSCs were lentivirally transduced to express Neurogenin 2 with a tetracycline regulatable system pLVX-UbC-rtTA-Ngn2:2A:EGFP (Addgene, #127288), subsequently cultures underwent puromycin selection (0.5 μg ml^−1^; Gibco) to enrich for transduced cells. For neuronal conversion, cells were transferred to PluriPro Matrix (Cell Guidance Systems) coated plates as a monolayer in PluriPro media (Cell Guidance Systems) with 10 μM ROCK inhibitor and were induced 24 hours after plating with 2 μg ml^−1^ doxycycline (Sigma) for two days. On day 3, cells were transitioned to laminin- and poly-L-lysine-coated plates in neuronal maturation media. Neuronal maturation media consisted of DMEM/F12 1:1 mix with Neurobasal (Gibco both), 1x B27 supplement (Gibco), 1 μg ml^−1^ laminin (Life Technologies), 2 μg ml^−1^ doxycycline (Stem Cell Tech), 0.5 mM dbCAMP (Stem Cell Tech), 20 ng ml^−1^ GDNF (Stem Cell Tech) and 20 ng ml^−1^ BDNF (Stem Cell Tech). On days 4-5, cells were also supplemented with 10 μM Ara C (Sigma) to remove proliferating cells. Neurons were cultured for a total of 14-21 days.

#### Fluorescence activated cell sorting (FACS)

For neurons, cells were dissociated when mature with Tryple (Gibco) and sorted based on eGFP expression with Zombie Violet (Biolegend) for live/dead discrimination. For microglia, cells were mechanically dislodged from Matrigel plates and treated with FC receptor block (1:20, Biolegend). Cells were then stained with the following 6 antibodies, all at 1:30 dilution and all from Biolegend: CD64-APC, CX3CR1 PCP-Cy 5.5, CD14-488, CD11b-PE, HLADR PC-Cy7, CD45-APC-CY7 for one hour. Cells were then washed and incubated in Zombie Violet (Biolegend) for live/dead discrimination. Controls consisted of cells incubated with a combination of appropriate isotypes for each antibody (Biolegend). Flow cytometry was performed on a BD InFlux Cytometer (Becton-Dickinson).

#### Immunocytochemistry

Cell cultures were fixed with 4% paraformaldehyde solution for 15 mins at room temperature. Antigen blocking and cell permeabilization were performed using 10% horse serum and 0.1% Triton X-100 in PBS for 1 hour at room temperature. Primary antibodies were incubated in 10% horse serum overnight at 4°C, and secondary antibodies (1:250, Jackson Laboratories) were incubated in the same solution for 1 hour at room temperature. The cells were counterstained with DAPI for nuclei detection. The following primary antibodies were used: rabbit anti-S100β (1:100; Abcam), chicken anti-GFAP monoclonal (1:1000; Abcam), mouse βIII Tubulin (1:500, Biolegend), chicken Map2 a+b (1:1000, Abcam), goat Iba-1 (1:1000, Novus), rabbit PU.1 (1:200, Cell Signaling), mouse TRA 1-60 (1:200, EMD Millipore), mouse TRA 1-81 (1:200, EMD Millipore), goat Nanog (1:200, R&D Systems), and rabbit SOX2 (1:500, Cell Signaling).

#### Western blotting

Western blotting was performed using standard procedures. The following antibodies were utilized: mouse GAPDH (1:10,000; Fitzgerald) and rabbit monoclonal BIN1 (1:5,000; Abcam). All secondary antibodies were purchased from Jackson ImmunoResearch, mouse or rabbit HRP (1:5,000).

#### Imaging

Imaging data were captured and processed using a confocal microscope (Zeiss LSM780). Image processing was performed using Zen software (Zeiss). Western image quantification was performed using ImageJ (NIH). Flow cytometry images were processed using FloJo Software (FlowJo).

#### Data analysis

##### Data preprocessing

FASTQ-files for ATAC-seq and ChIP-seq were processed with the official ENCODE ATAC-seq (https://github.com/ENCODE-DCC/atac-seq-pipeline) and ChIP-seq (https://github.com/ENCODE-DCC/chip-seq-pipeline2) pipelines, respectively. Both pipelines were installed with a local anaconda environment. The Build_genome_data scripts were run to build the local hg19 reference genome and other dependencies. These pipelines execute a number of programs to process and align sequencing reads and to generate quality control statistics (Table S2). In brief, FASTQ-files were trimmed with cutadapt (Version 1.9.1) and aligned with Bowtie2 (*90*) to hg19. SAMtools (*91*) (Version 1.2), MarkDuplicates (Picard Version 1.126), and bedtools (*92*) (Version 2.26) were used to filter and clean data post alignment. MACS2 (Version 2.1.1) was used for peak calling (*93*). The ATAC-seq pipeline also runs IDR (*94*) to detect peaks with high reproducibility. The optimal peak sets that came out of these pipelines were used for downstream analysis.

#### RNA-seq data processing

FASTQ-files were mapped to the UCSC genome build hg19 with STAR (Version 2.5.3) using default parameters (*95*), and converted into homer tag directories (*96*). The HOMER function “analyzeRepeats” was used to quantify raw reads with the parameters “-raw -count exons -condenseGenes” and normalized reads, as either counts per million (CPM) values using “-cpm -count exons -condenseGenes”, or as transcripts per million (TPM) values using “-tpm -count exons -condenseGenes.” The base-2 logarithm of the TPM/CPM values was taken after adding a pseudocount of 1 TPM/CPM to each gene. For the iPSC experiments, differential analysis was done with the HOMER function “getDiffExpression” with the parameters “-repeats -DESeq2 -AvsA” with an FDR < 0.05 and log_2_ fold change > 1.

#### Enhancer and promoter calling

For each cell type, active promoters were identified by H3K4me3 peaks that overlap H3K27ac within 2,000 bp to a nearest transcription start site (TSS). Active enhancers were identified by H3K27ac peaks that were outside of H3K4me3 peaks.

#### LDSC regression analysis

To determine whether specific brain cell type annotations were enriched for heritability of specific neurological disorders, psychiatric disorders and neurobehavioral traits, stratified LD score regression was applied (*27, 97*). Table S4 contains an overview of all the GWAS datasets that were used. The cell type-specific enhancer and promoter peak sets were tested for enrichment of heritability while controlling for the full baseline model. The corresponding P-values were FDR multiple testing corrected for the number of GWAS studies and number of cell type-specific annotations.

#### Quantification of ChIP-seq and ATAC-seq datasets

Homer (*96*) (Version v4.9.1) was used to make tag directories of H3K27ac and H3K4me3 ChIP-seq, and ATAC-seq datasets belonging to each individual sample and for all combined samples per cell type. Quantification of H3K27ac and H3K4me3 ChIP-seq and ATAC-seq signal at peaks merged for all cell types was performed using the Homer annotatePeaks function. The Homer annotatePeaks function was used to quantify H3K27ac reads around TSSs (+/- 2000 bp) for differential gene activation analysis.

#### Differential analyses

To identify gene promoters that showed increased activation in a given cell type, we performed a differential analysis on the H3K27ac at TSSs with Limma Voom (*98, 99*) (Version 3.34.9). A linear model was set up with four contrasts: each cell type versus the average of the other cell types. Genes that were at least 4-fold induced with an FDR corrected P-value below 0.01 were combined and used to generate cell type-specific gene signatures. For genes with multiple TSSs, the TSS quantification with the highest number of reads was selected as a proxy for gene expression. These cell type-specific gene activation signatures were subsequently used for metascape enrichment analyses.

#### Correlations across assays

RNA-seq FASTQ-files from different cell types of the brain were downloaded from SRA and processed as described above (*21*). Hippocampal, fetal, bulk whole-cortex, endothelial, and tumor-derived tissues were excluded. Spearman correlations between log_2_(CPM) of RNA-seq with H3K27ac and H3K4me3 ChIP-seq and ATAC-seq around TSSs were calculated for all replicate combinations. Correlations were classified as ’cell type correlations’ when RNA-seq from one cell type was compared with either H3K27ac ChIP-seq, H3K4me3 ChIP-seq or ATAC-seq from the same cell type. Correlations were classified as ‘aspecific correlations’ when RNA-seq from one cell type was compared with either H3K27ac ChIP-seq, H3K4me3 ChIP-seq or ATAC-seq from a different cell type. The significance of the difference of these distributions of correlations was calculated with Kruskall-Wallis between group test.

#### Cell type purity analyses

Gene signatures for individual brain cell types were collected from psychEncode (*25*). For the ‘inhibitory neurons’ and ‘excitatory neurons’, all the gene signature sets corresponding to excitatory (Ex1-8) and inhibitory (In1-8) neurons were combined. The distributions of H3K27ac log_2_(CPM) values at TSSs for each pooled tag directory were compared between cell types using Kruskall-Wallis between group test. The Dunn post-hoc test was applied to calculate the significance of each pairwise combination with a Bonferroni multiple testing correction. The distributions were visualized as a violin plot, with asterisk on the bottom that correspond to the Kruskall-Wallis between group p-values, and lines and asterisk between individual significant pairwise differences at the top. *** = p < 1e-8; ** = p < 5e-5, * p < 5e-2.

#### Motif analysis

To find transcription factors that were associated with each cell type, we performed a de novo motif analysis on accessible chromatin peaks within 500 bp of enhancers. Motif enrichments were performed with HOMER function findMotifsGenome. The output report links de novo motifs to known motifs and gives a motif matching score. Known motifs that matched to the de novo motif with a score higher than 0.7 were linked to transcription factors based on the human transcription factor catalog (*44*). The human transcription factor catalog was used to determine transcription factors that were associated with disease based upon proximal GWAS signal (*44*). N.C., a board-certified pediatrician and pediatric intensivist, manually curated this list for conditions that are related to the brain.

#### Fine mapping

Fine mapping was performed with PAINTOR Version 3 (*62*) and CAVIAR Version 2.2 (*61*). Linkage Disequilibrium (LD) values between all SNPs in significant AD loci (Stage 1)(*28*) were calculated with PAINTOR function CalcLD_1KG_VCF, with the 1000G European reference cohort. Loci were extended to minimally encompass 20,000 bp. The microglia H3K27ac peak file was used to guide fine mapping with PAINTOR, with an enumerate value of 1. CAVIAR was run on the same LD data with max causal-variants value of 1.

#### Gene Ontology (GO) Enrichment analysis

Metascape (http://metascape.org), with input species set to Homo sapiens, was used to identify GO terms and pathways that were overrepresented in particular gene sets (*100*) such as cell type-specific H3K27ac ChIP-seq at TSSs and cell type-specific PLAC-seq-linked genes. The resulting FDR corrected P-values were depicted as -log10(q) bar plots.

#### String analysis

The string database of protein-protein interactions (*101*) (https://string-db.org) was used to analyze the extended AD-risk gene set, defined as gene promoters with AD-risk SNPs and/or PLAC-linked to AD-risk SNPs. The string database was used to determine whether the extended AD-risk gene set was enriched for protein-coding genes that are in a predicted protein-protein interaction network for microglia, neurons and oligodendrocytes.

#### PLAC-seq processing

FASTQ-files were trimmed with TrimGalore (version 0.4.5), which is a wrapper around Cutadapt (version 1.1.5). PLAC-seq data was processed with MAPS (*58*) (downloaded from github on October 4^th^ 2018). In short, MAPS aligned the trimmed FASTQ-files with BWA to a hg19 reference genome for the R1 and the R2 reads separately. Aligned reads were paired and valid pairs were used for downstream application. MAPS normalizes chromatin contact frequencies to detect significant chromatin interactions anchored at genomic regions at the merged H3K4me3 peaks (for Olig2, NeuN and PU.1) to identify long-range chromatin interactions at 5,000 bp resolution. Significant interactions were identified with FDR corrected P-value cutoff of 0.01. MAPS was run on all samples individually and on a merge of the samples that belong to the same cell type. The significant interactions for all samples belonging to each cell type were merged in a general interaction set, and for each individual sample these interactions were quantified. The quantified matrix of all significant interactions for all cell types was used as input for Limma differential analysis. A linear model was fit, with three pairwise contrasts. Interactions that were significantly increased in both comparisons of each cell type vs other cell types (FDR < 0.05, and log_2_ FC>1) were taken as cell type-specific increased interactions.

To classify the functional elements belonging to the loops and calculate the proportions, the interactions were classified in a hierarchical fashion for genomic annotations belonging to the same cell type. First, it was determined if both bins contained an active promoter (subsequently classified as a promoter-promoter interaction). If not, it was determined if the loops contained an active promoter in one bin and an enhancer in the other bin (enhancer-promoter interaction), or if the loop links a promoter to an ATAC-seq peak (promoter-ATAC interaction), or if the promoter was not linked to any peaks (promoter to none). If there were no active promoters within either bin of the interactions, we determined if there was a TSS-distal H3K4me3 peak in either bin (classified as distal H3k4me3 bound interaction). Interactions without classifications were assigned ‘unclear’.

#### UCSC genome browser

For ATAC-seq and ChIP-seq signals, the HOMER function makeMultiWigHub was used to generate UCSC track hubs from HOMER tag directories. For PLAC-seq interactions, significant MAPS interactions were converted into bigbed with UCSCtool bedToBigBed (*102*) with type = bed5+13 for the merged samples belonging to each cell type. UCSC genome browser (*103*) was used to generate genome browser shots of loci of interest.

#### Super-enhancers

Super-enhancers were identified using HOMER’s findPeaks command on the pools of H3K27ac reads with pooled input reads as background with the parameters “-style super -L 1”.

#### Heatmaps

Heatmaps were made in R with pheatmap (Version 1.0.8) (https://CRAN.R-project.org/package=pheatmap).

#### Circos plot

The Circos plot was generated with BioCircos (*104*) (0.3.3) R Package.

#### Chow Ruskey plots

Chow-Ruskey plots were made with R package Vennerable (https://github.com/js229/Vennerable Version 3.1.0.9).

#### Additional visualizations

Violin plots, bar plots and circle plots were made with Seaborn (Version 0.8.1) and matplotlib (*105*) (Version 2.2.2).

#### Python and R environments

The Jupyter notebook was used for all analyses from Anaconda (Conda version 4.6.9) with python (version 3.6.5) and R (version 3.4.3). Statistical analyses were performed with Scipy (version 1.1.0) and Numpy (version 1.1.5) with Pandas libraries (version 0.23.4). Post-hoc tests were calculated with scikit-posthocs library (Version 0.6.1).

### Supplemental Figures

**Fig. S1.**
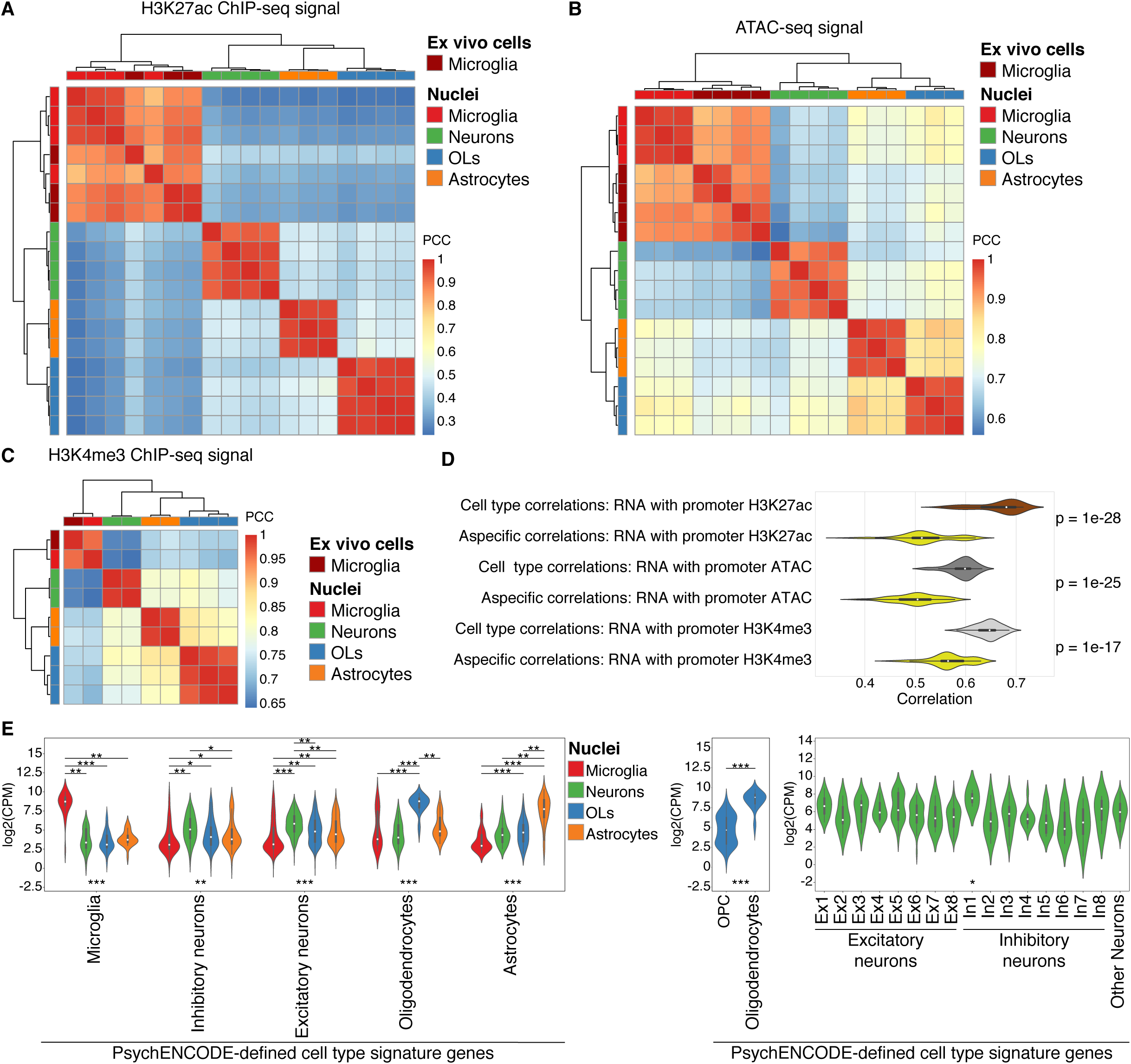
Cell type-specific isolation of nuclei populations from the brain. (A) Heatmap of Pearson’s correlation of H3K27ac ChIP-seq log_2_(CPM) in ex vivo microglia and nuclei of microglia, neurons, astrocytes and oligodendrocytes. (B) Heatmap of Pearson’s correlation of ATAC-seq log_2_(CPM) in ex vivo microglia and nuclei of microglia, neurons, astrocytes and oligodendrocytes. (C) Heatmap of Pearson’s correlation of H3K4me3 ChIP-seq log_2_(CPM) in ex vivo microglia and in nuclei of microglia, neurons, astrocytes and oligodendrocytes. (D) Violin plots depicting the distribution of Pearson’s correlations of RNA-seq log_2_(CPM) values with ATAC, H3K27ac and H3K4me3 log_2_(CPM) values at gene promoters between replicates of the same cell types (cell type correlations) and between replicates of different cell types (aspecific correlations). (E) Violin plots (left) of brain nuclei populations showing log_2_(CPM) values of H3K27ac signal at the promoters of cell type signature genes for microglia, excitatory and inhibitory neurons, astrocytes and oligodendrocytes as defined by PsychEncode (*25*). Violin plot (middle) of oligodendrocyte nuclei showing log_2_(CPM) values of H3K27ac signal at the promoters of cell type signature genes for OPCs and oligodendrocytes. Violin plot (right) of neuronal nuclei showing log_2_(CPM) values of H3K27ac signal at the promoters of cell type signature genes for subclasses of excitatory and inhibitory neurons. Kruskal-Wallis-between group test and Dunn post-hoc test; bottom asterisks shows Kruskall-Wallis between group p-values; upper asterisks show pairwise significant p-values; *** = p < 1e-8; ** = p < 5e-5, * p < 5e-2. PCC, Pearson’s correlation coefficient.

**Fig. S2.**
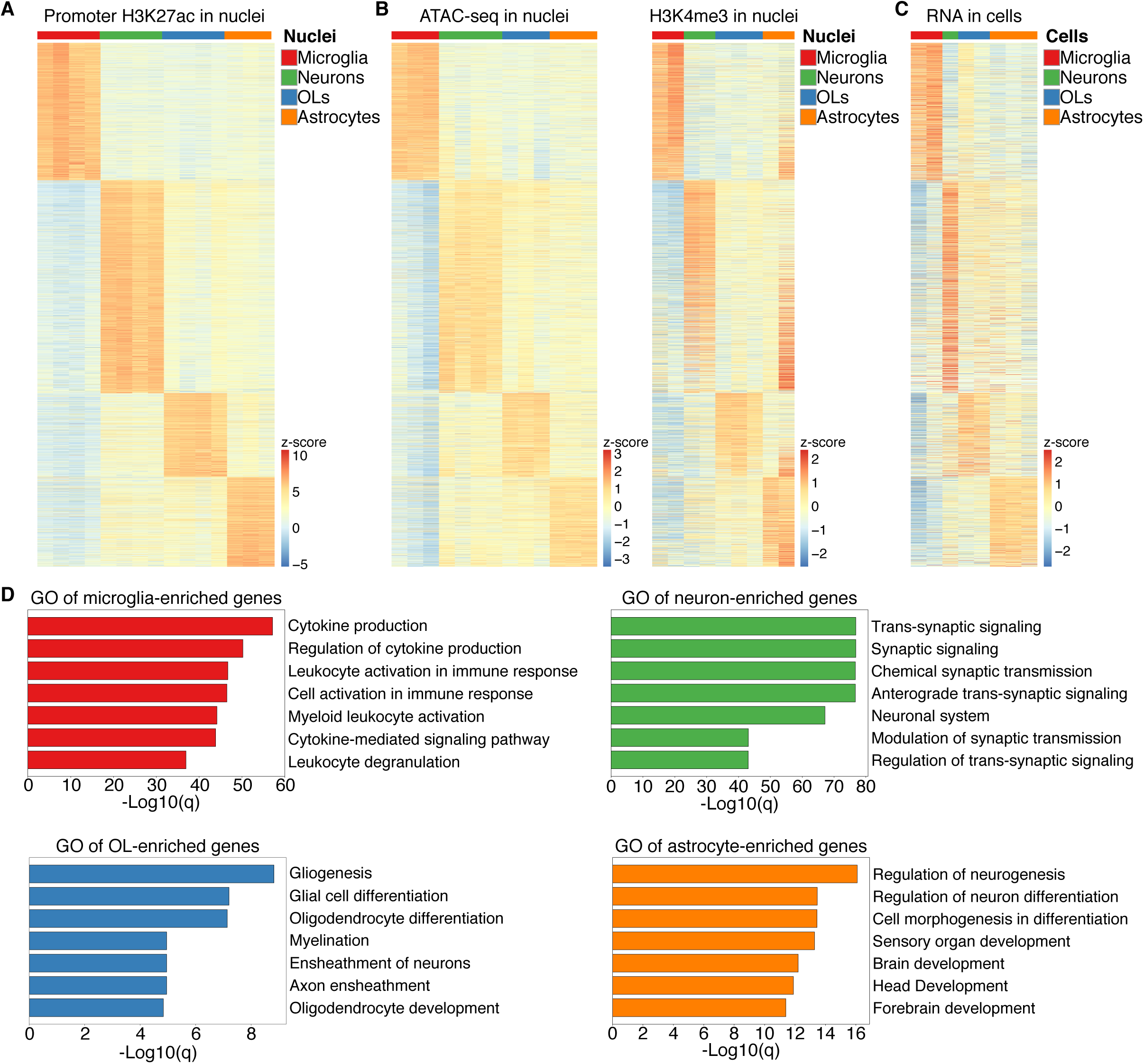
Cell type-specific isolation of nuclei populations from the brain. (A) Heatmap of z-score normalized log_2_(CPM) values of promoter H3K27ac signal for genes with cell type-enriched promoter H3K27ac versus other brain cell types (FDR < 0.01 and logFC > 2). (B) Heatmap of z-score normalized log_2_(CPM) of ATAC-seq and H3K4me3 signal at gene promoters identified and shown in the same order as the H3K27ac heatmap in (A). (C) Heatmap of z-score normalized log_2_(CPM) values of cell type-specific RNA-seq (*21*) at genes corresponding to promoters identified and shown in the same order as the H3K27ac heatmap in (A). (D) Metascape analyses of genes with differentially active promoters as identified in (A) depicted as –log_10_(q)-values.

**Fig. S3.**
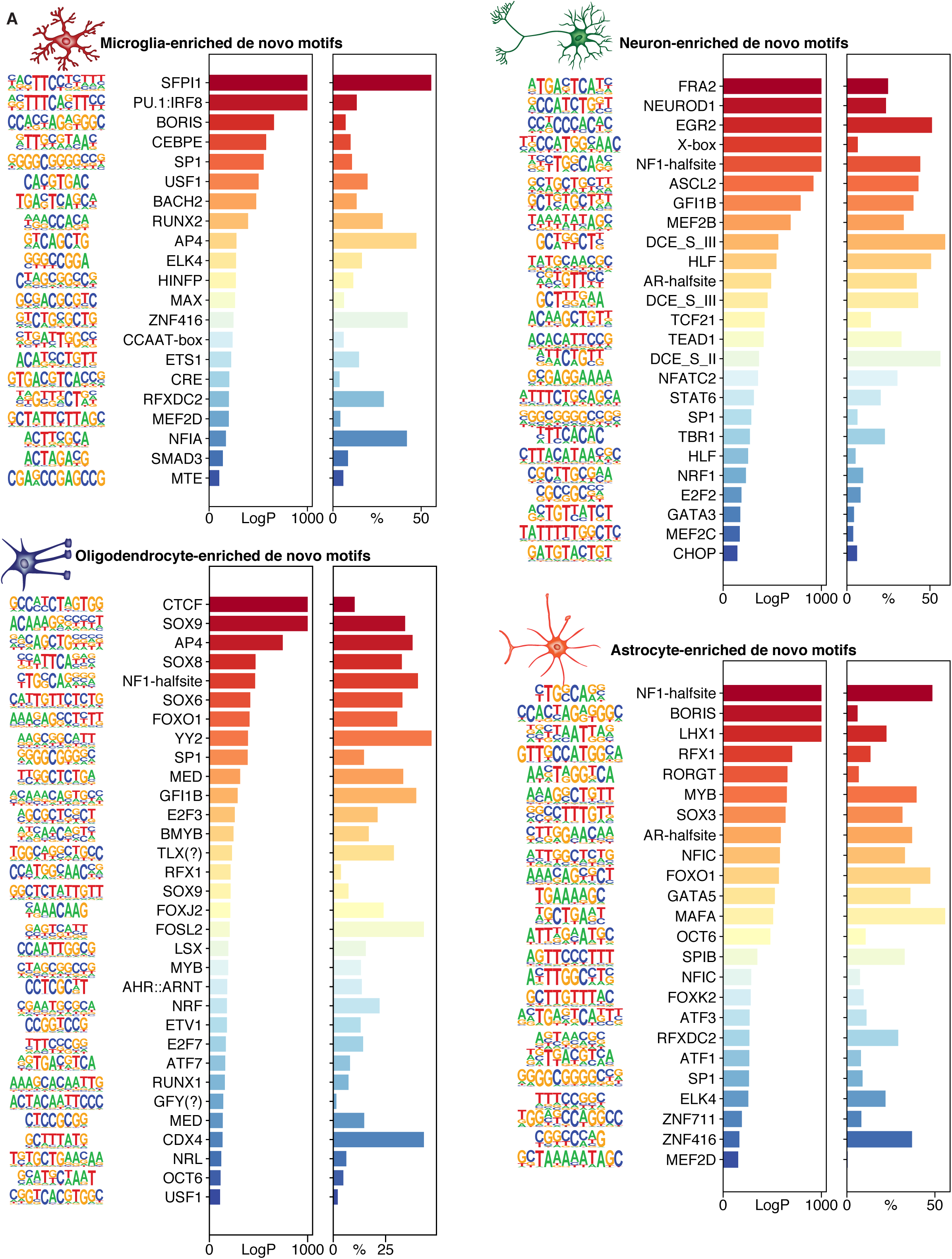
Enrichment of de novo motifs at brain cell type enhancers. Significant de novo motifs identified by HOMER in regions of open chromatin at active enhancers in nuclei of microglia, neurons, astrocytes and oligodendrocytes. For each de novo motif is displayed the –log_10_(P) values and the percentage of peaks containing the motif.

**Fig. S4.**
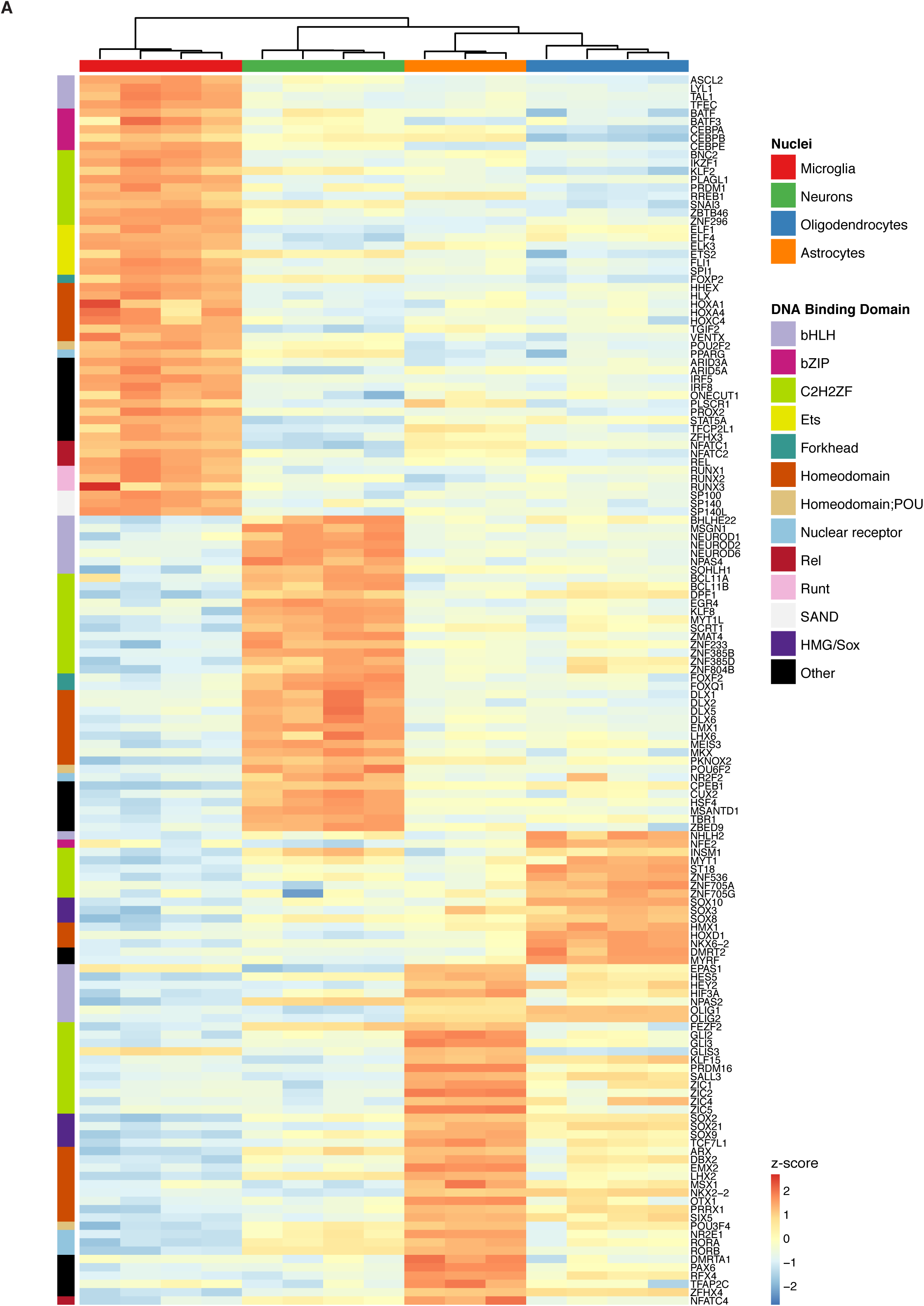
Differential promoter H3K27 acetylation of human transcription factor genes in cell types of the brain. Heatmap of z-score normalized log_2_(CPM) values of H3K27ac at promoters of the top 148 transcription factors with cell type differential H3K27ac signal (FDR < 0.01 and logFC > 3) in microglia, neurons, astrocytes and oligodendrocytes using a compilation of 1639 human transcription factors (*44*).

**Fig S5.**
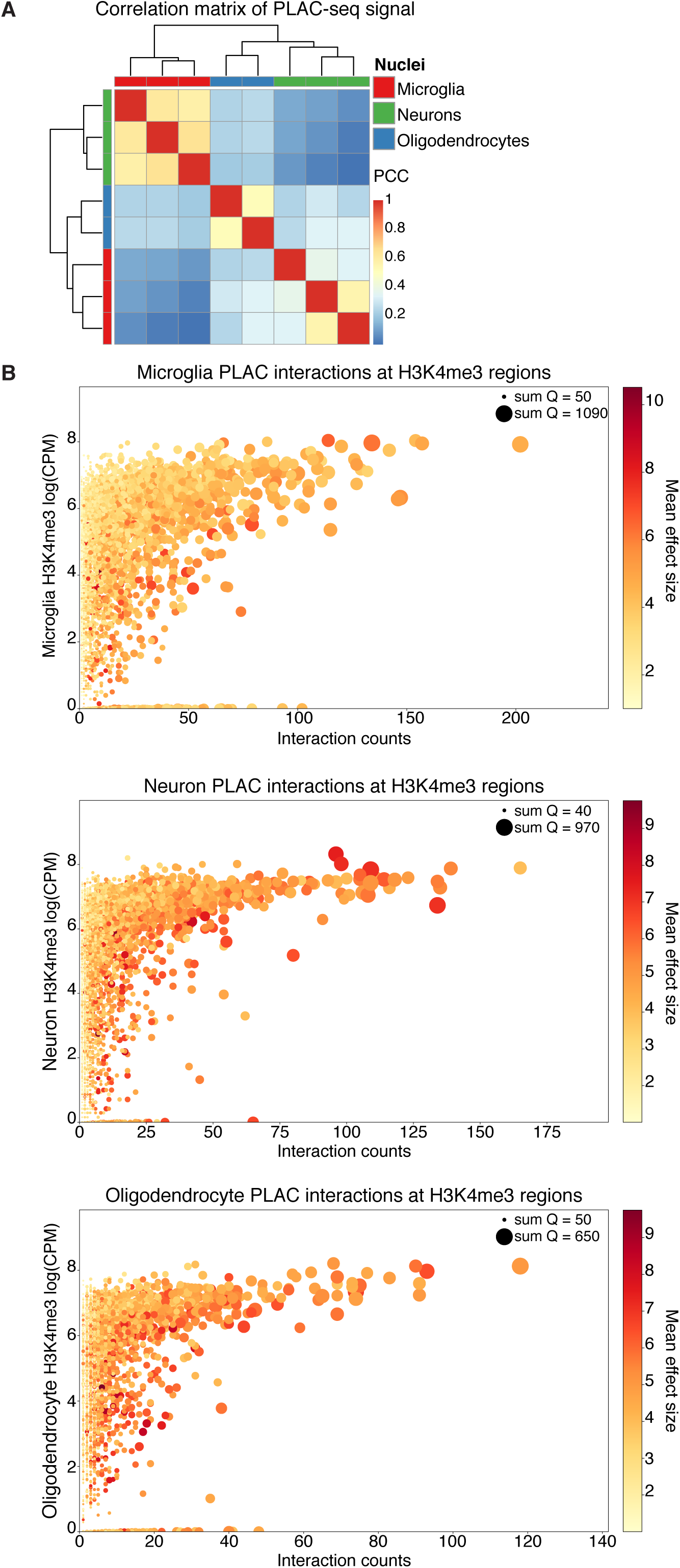
Chromatin interactions in cell types of the brain. (A) Pearson’s correlation heatmap of PLAC-seq log_2_(CPM) values for microglia, neuron and oligodendrocyte replicates; PCC, Pearson’s correlation coefficient. (B) Scatterplots of H3K4me3 log_2_(CPM) values (y-axis) and PLAC-seq interaction counts (x-axis) for microglia (top), neurons (middle) and oligodendrocytes (bottom). Each data point represents a gene, the mean effect size is depicted in the color gradient of each data point and the cumulative log10(q-values) is depicted as the size of each data point.

**Fig S6.**
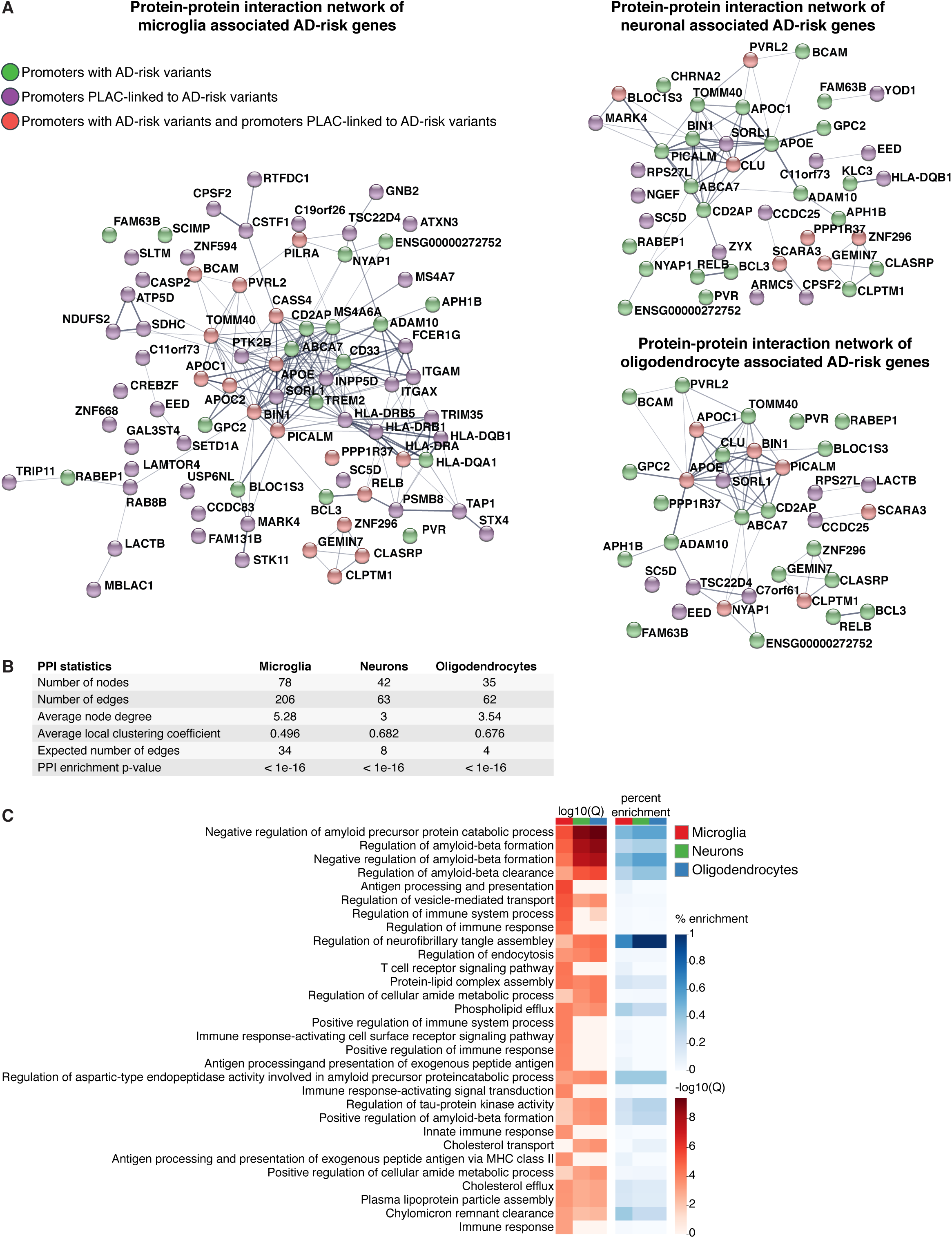
Chromatin interactions at AD loci for cell types of the brain. (A) STRING Protein-protein interaction (PPI) network of genes that have either (1) promoters with AD-risk variants (green), (2) promoters PLAC-linked to AD-risk variants (purple) or (3) promoters with AD-risk variants and PLAC-linked to AD-risk variants (green/purple) identified in microglia (left), neurons (right, top) and oligodendrocytes (right, bottom). Line thickness indicates level of confidence. (B) Table of protein-protein interaction summary statistics for microglia, neurons and oligodendrocytes. (C) Heatmaps of STRING gene ontology analyses of AD-risk genes identified in microglia, neurons and oligodendrocytes showing -log_10_(q) values (red) and percent enrichment (blue).

**Fig. S7.**
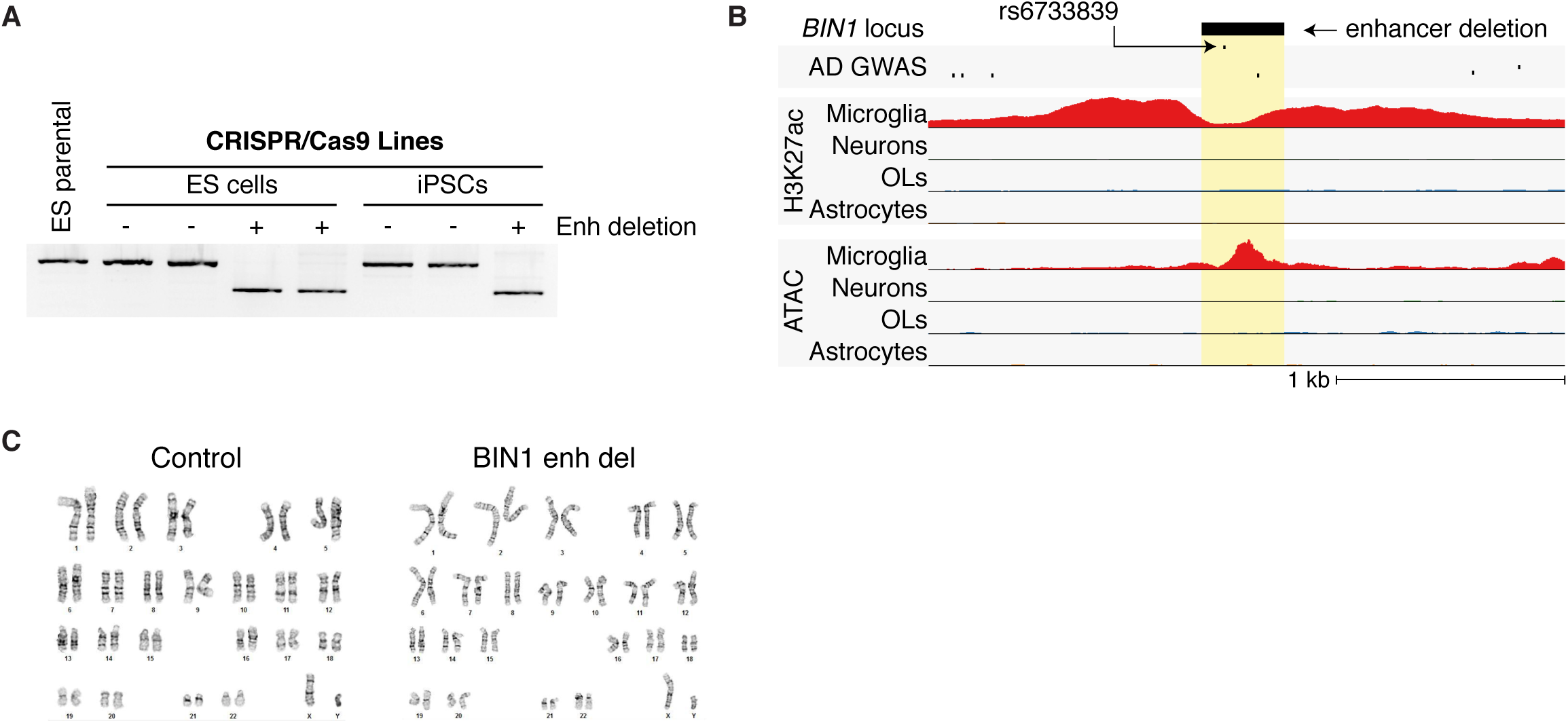
Deletion of a microglia enhancer harboring a lead AD-risk variant upstream of the *BIN1* locus. (A) PCR amplification of the *BIN1* enhancer region in control and *BIN1*^enh_del^ PSCs delineating the 363bp enhancer deletion. (B) UCSC genome browser visualization of an enhancer at the *BIN1* locus that overlaps the rs6733839 AD-risk variant shows ATAC-seq and H3K27ac ChIP-seq signal restricted to microglia. A 363 bp CRISPR/Cas9-deleted region is shown as a black bar (highlighted yellow). (C) All control and *BIN1*^enh_del^ PSCs retained a normal karyotype.

**Fig. S8.**
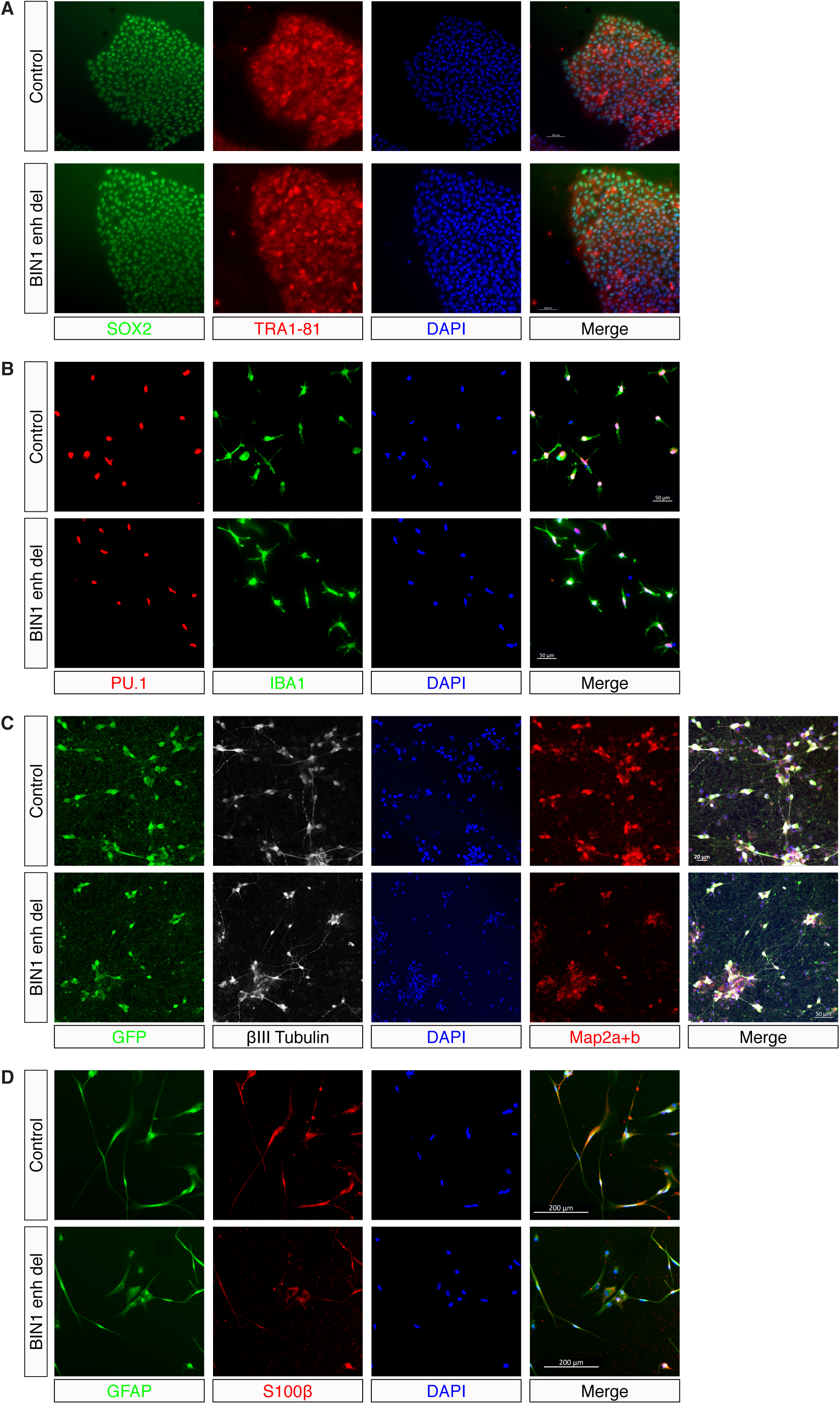
Control and *BIN1*^enh_del^ PSCs and derived microglia, neurons and astrocytes express typical cell lineage markers. Representative immunohistochemistry images of PSCs (A), microglia (B), neurons (C) and astrocytes (D) in control and *BIN1*^enh_del^ lines stained for the indicated cell lineage markers.

**Fig. S9.**
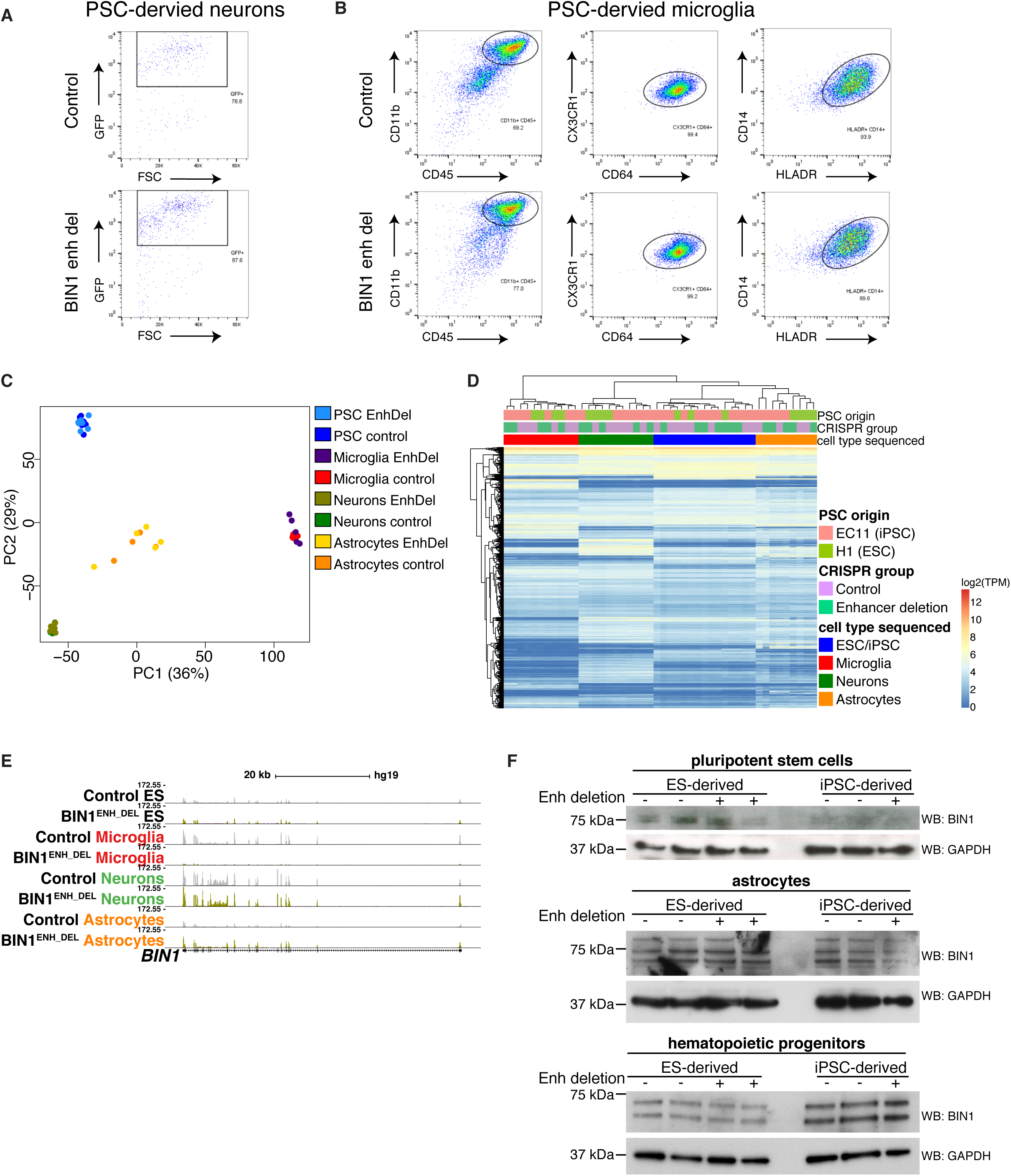
Deletion of a microglia enhancer harboring a lead AD-risk variant affects *BIN1* expression in PSC-derived microglia (A) Control and *BIN1*^enh_del^ neurons were sorted for expression of Neurogenin2 with 2A peptide-linked eGFP (B) Control and *BIN1*^enh_del^ microglia were sorted for expression of CD11B^high^, CD45^intermdetiate^, CX3CR1^+^, CD64^+^, CD14^+^, and HLADR^+^. (C) Principal component analysis of RNA-seq log_2_(TPM) for control and *BIN1*^enh_del^ PSCs, microglia, neurons and astrocytes. (D) Heatmap of RNA-seq log_2_(TPM) for control and *BIN1*^enh_del^ PSCs, microglia, neurons and astrocytes. (E) UCSC browser visualization of RNA-seq tag counts at the *BIN1* locus for control and *BIN1*^enh_del^ PSCs, microglia, neurons and astrocytes. (F) Western blot of BIN1 and GAPDH in control and *BIN1*^enh_del^ PSCs (top), and derived astrocytes (middle) and hematopoietic progenitors (bottom).

**Table S1** Patient information and assays

Table summarizing patient information including age, sex, ethnicity, brain region and pathology diagnosis. Table summarizes isolation method for either ex vivo cells or nuclei, cell type of origin and assays performed.

**Table S2** ENCODE quality control statistics

Table summarizing quality control statistics generated from the ENCODE pipelines used to process ATAC-seq, H3K27ac ChIP-seq and H3K4me3 ChIP-seq datasets for replicates of each cell type (see tabs). Table includes IDR statistics generated from the ENCODE pipeline used to process ATAC-seq datasets for replicates of each cell type (see tabs).

**Table S3** Differential promoter H3K27ac analysis

Table with results of differential H3K27ac around TSS. Columns represent log_2_FC of each cell type versus the other cell types and FDR multiple testing corrected p-values. The Boolean ‘is_TF’ column represents whether a gene is annotated as a transcription factor.

**Table S4** Summary of GWAS studies

Table summarizing GWAS studies used for LDSC regression analysis in this study.

**Table S5** Cell type-specific transcription factors and disease association

Table summarizing a subset of cell type-specific transcription factors that were paired with enhancer motifs of the corresponding cell type. Table includes information on diseases associated with transcription factors based upon proximal GWAS signal (*44*), which has been subset for conditions that are related to the brain.

**Table S6** PLAC seq promoter interactome map

Promoter centric interactome map that links promoters active in one or multiple cell types to distal regulatory regions that contain one or multiple active promoters and or active enhancers.

**Table S7** PLAC seq interactions at super enhancers

Table summarizing PLAC-seq promoter interactions with super enhancers for each cell type, including information of GWAS variants localized to PLAC-seq linked super-enhancers. The Boolean columns indicate whether the super-enhancer overlaps with GWAS variants for neurodegenerative diseases (AD, amyotrophic lateral sclerosis, epilepsy and autism), psychiatric conditions (major depressive disorder, schizophrenia), and neurobehavioral traits (intelligence, cognitive function, risk behavior, neuroticism and insomnia). The numerical columns indicate the number of GWAS variants that overlap within a super-enhancer for each condition.

**Table S8** Differential gene expression of *BIN1* enhancer deletion lines

Table provides DESeq2 output as rlog transformed data of RNA-seq data for PSCs and PSC-derived microglia, neurons and astrocytes for each gene. Table includes log_2_ fold change values, p-values and adjusted p-values for each gene between control and *BIN1* enhancer deletion groups for PSCs and PSC-derived microglia, neurons and astrocytes.

